# Building Trust in Deep Learning-based Immune Response Predictors with Interpretable Explanations

**DOI:** 10.1101/2023.05.02.539109

**Authors:** Piyush Borole, Ajitha Rajan

## Abstract

The ability to predict whether a peptide will get presented on Major Histocompatibility Complex (MHC) class I molecules has profound implications in designing vaccines. Numerous deep learning-based predictors for peptide presentation on MHC class I molecules exist with high levels of accuracy. However, these MHC class I predictors are treated as black-box functions, providing little insight into their decision making. To build turst in these predictors, it is crucial to understand the rationale behind their decisions with human-interpretable explanations. We present MHCXAI, eXplainable AI (XAI) techniques to help interpret the outputs from MHC class I predictors in terms of input peptide features. In our experiments, we explain the outputs of four state-of-the-art MHC class I predictors over a large dataset of peptides and MHC alleles. Additionally, we evaluate the reliability of the explanations by comparing against ground truth and checking their robustness. MHCXAI seeks to increase understanding of deep learning-based predictors in the immune response domain and build trust with validated explanations

## 1 Introduction

The Major Histocompatibility Complex (MHC) class I pathway supports the detection of cancer and viruses by the immune system. It presents parts of protein (peptides) from inside a cell on to the membrane surface enabling visiting immune cells that detect non-self peptides to terminate the cell. The ability to predict whether a peptide will get presented on MHC class I molecules is a key component in vaccine design since it helps determine if the vaccine can activate the immune system to destroy the invading pathogen. Numerous deep learning (DL)-based models for predicting peptide presentation on MHC class I molecules have emerged, demonstrating high prediction accuracies in predicting presented peptides. The deep learning models in the MHC-I predictor literature have a wide variety of architectures - Multilayer perceptron (NetMHCpan-4.1 [43]), Convolutional Neural Networks (MHCfovea [29], ConvMHC [19]), Transformers (TransPHLA [11], ACME [23]), Gated Recurrent Unit neural networks (MHCSeqNet [40]), etc. However, owing to the inherent inscrutability of deep learning models, it is difficult to understand and interpret predictor performance for the peptide-MHC allele instances and rationalize the differences observed between the predictors. Explainable Artificial Intelligence (XAI) techniques have been proposed in recent years to help explain and understand the output of deep learning models used in image classification [44, 60, 51, 17, 49, 52, 48] and NLP classification tasks [18, 1, 32, 13, 34]. There is, however, limited work on explainability and interpretability for deep learning-based MHC class I predictors.

Related work for MHC class 1 predictors largely focus on global explainability, that is trends and explanations observed across the whole input dataset, rather than local explainability focusing on individual input instances. ACME and TransPHLA use attention scores to provide both global explanation and explanation for just one input (Instance-based or local explanation). However, the use of attention scores as an explanation is not reliable [28] and is architecture specific, making it unusable for most MHC class I predictors that use other architectures as Convolutional Neural Networks or Multilayer perceptron. PoSHAP, proposed by [15] in 2022, is related to the interpretations that we use. However, PoSHAP in their contribution only consider the Long Short-Term Memory (LSTM) deep learning architecture and focus their analysis on producing global, rather than local, explanations. Our work focuses on post-hoc explanations, i.e., explanations for an existing model that has been previously trained, which we treat as a black-box system. Such post-hoc explanations are widely applicable as they can be used over predictors whose internal structure is not known.

As a first contribution in this article, we use two popular XAI techniques, Locally Interpretable Model Agnostic Explanations (LIME) [44] and SHapley Additive exPlanation (SHAP) [33], from the image classification domain to *interpret the outputs from MHC class I predictors*. Both XAI techniques are model agnostic and can be applied to any deep learning-based MHC class I predictor, irrespective of architecture. LIME perturbs (or mutates) the input to create samples for training a simpler model such as linear regression. These simpler models approximate the complex model in the input locality and its features can be interpreted as explanations. SHAP also uses mutation of the input but employs a concept from game theory namely ‘Shapley values’ to determine contribution of each input feature to the final prediction.

We use four state of the art MHC class I predictors with different deep learning architectures and generate explanations for their outputs with both XAI techniques. The explanations for MHC class I predictors highlight important regions of the input peptide used in the output prediction which can be used for model debugging and interpretability. Interpretable explanations are key to building trust in deep learning models, in line with safety regulations such as the recently proposed EU AI act https://artificialintelligenceact.eu/the-act/ that requires explanations to help users better understand the decisions made by AI systems.

Although current XAI techniques for deep learning architectures have created a step change in providing reasons for predicted results, the question of whether the explanations themselves can be trusted has been largely ignored. Some recent studies [41, 8, 20] have demonstrated the limitations of current XAI techniques. For instance, [41] applied three different XAI techniques on a CNN-based breast cancer classification model and found the techniques disagreed on the input features used for the predicted output and in some cases picked background regions that did not include the breast or the tumour as explanations. Literature on evaluating the reliability of XAI techniques is still in its nascency and can be broadly divided into two branches - (1) Studies that assume the availability of expert annotated ground truth, maybe in the form of bounding boxes for images, to evaluate the accuracy of explanations [30, 61, 57, 62, 58, 7, 2, 21] and (2) research that uses the idea of removing relevant (or important) features detected by an XAI method and verifying the accuracy degradation of the retrained models [22, 39, 46, 59, 27, 6, 31, 10]. The first category requires human-annotated ground truth for evaluation while the second category incurs very high computational cost to verify accuracy degradation from retraining the models.

As our second contribution, we evaluate the *quality and reliability of explanations* generated by the two XAI techniques, LIME and SHAP. We do this in three different ways, (1) We check if the explanations match ground truth which in the ideal case is exact information on peptide positions involved in the binding with MHC allele. This information is, however, unavailable. Instead, a proxy ground truth that can be used is the binding residues identified by BAlaS [56, 26] that calculates the difference between the free-energy of binding of original bound complex and mutated bound complex where just one residue of ligand peptide is replaced with alanine [56, 26]. (2) We check the consistency of LIME or SHAP by measuring the extent to which the explanation for a given input peptide is similar across different MHC predictors, and (3) We assess the stability of generated explanations by measuring similarity of explanations for similar input peptides with a given MHC predictor. interaction free energy

In summary, the article makes the following contributions, as shown in Fig. 1,

**Figure 1:**
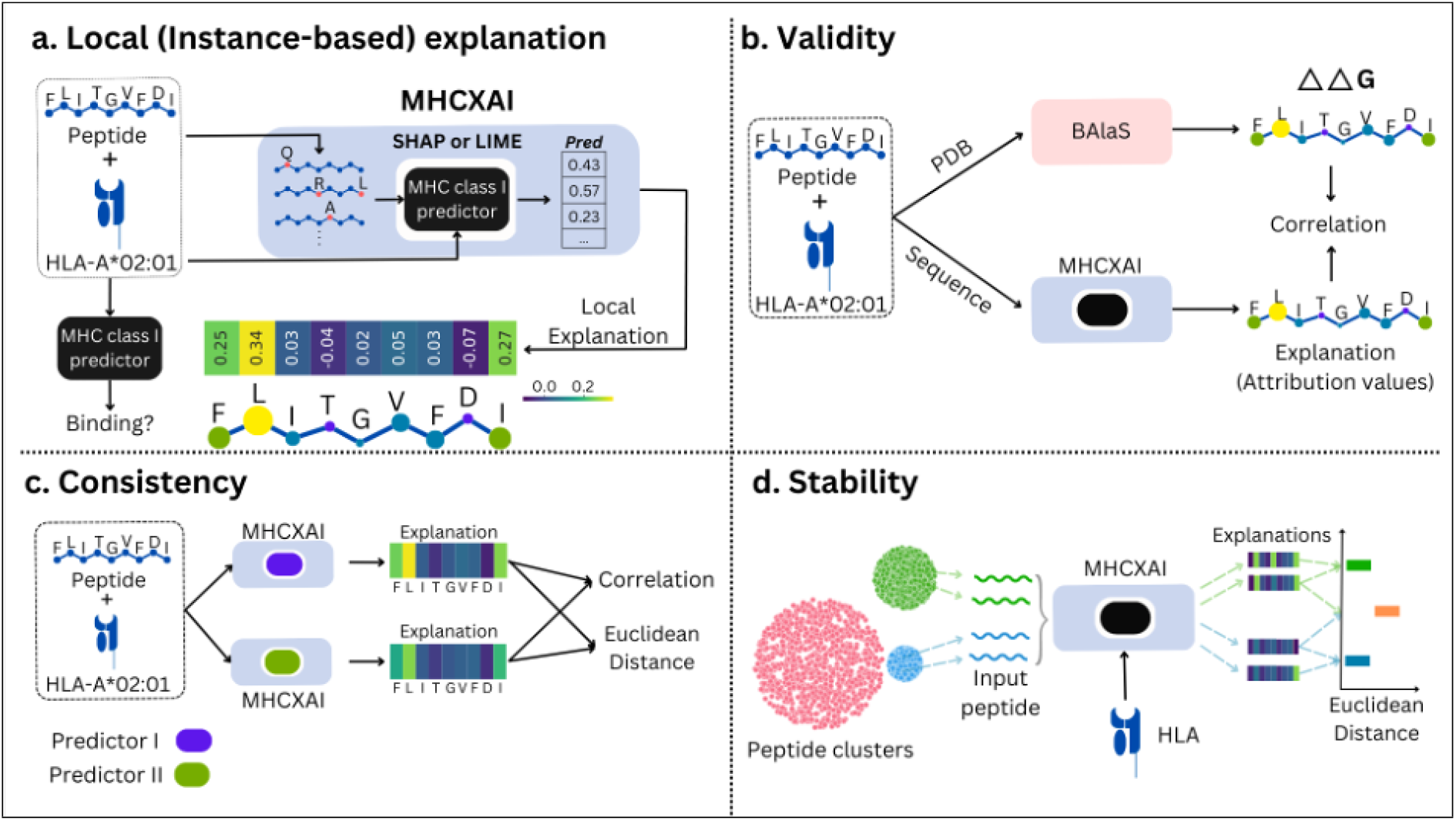
Explainable AI for MHC class I prediction. **a)** The framework that combines XAI techniques (SHAP or LIME) and MHC class I predictor is termed as `MHCXAI`. For generating explanations for input peptide, the peptide sequence is mutated (or perturbed) multiple times. Then, binding is predicted using MHC class I predictors for these mutated (or perturbed) sequence against the allele of interest. These inputs and their corresponding outputs are used to generate attribution values (explanation) for each peptide position. We visualize these values as heatmaps such that positions contributing positively to the positive class are highlighted with lighter color while negative contributions to the positive class are highlighted with darker colors. **b)** Validity of the generated explanation is tested by comparing them against important positions highlighted by BAlaS using the PDB structure of peptide-MHC allele bound complex. The quality of explanations are tested using XAI metrics `Consistency` **c** and `Stability` **d. c)** `Consistency` is the metric of XAI where it produces similar explanations for predictors with similar performances for a given input. We compare the two explanations using Pearson correlation coefficient and the Euclidean distance. **d)** `Stability` is the metric of XAI where it produces similar explanations for similar inputs (and similar prediction outcome) for a predictor. For every MHC allele, we cluster the binding peptides with GibbsCluster and generate explanations for them using LIME or SHAP. We compare the Euclidean distance between the explanations for peptides within a cluster (Intracluster)

1. A framework, `MHCXAI`, that provides novel instance-based explanations for MHC class 1 predictors using XAI techniques, LIME and SHAP (Fig. 1a). As part of this contribution, we evaluate four state-of-the-art (SOTA) MHC-I predictors over a large dataset, `MHCBench`, that we curate from several existing datasets

**Table 1:** Overlap between training peptides for MHC class I predictors and `MHC-Bench` dataset peptides
2. Assess the validity of the explanations generated from SHAP and LIME by comparing against BAlaS-based ground truth (Fig. 1b).
3. Assess quality of the explanations generated from two explainable AI techniques, shown in Fig. 1c and Fig. 1d, with respect to consistency and stability of explanations.

| **MHC class I predictor** | **# training peptides** | **# overlapping peptides** | **Overlap %** |
| --- | --- | --- | --- |
| MHCflurry | 190,095 | 95,743 | 4.29% |
| MHCfovea | 108,092 | 50,072 | 2.24% |
| NetMHCpan | 31,212 | 1,149 | 0.05% |
| TransPHLA | 1,184,191 | 289,605 | 12.97% |

## 2 Results

### 2.1 Evaluation of MHC class I predictors

Here, we report performance of four state-of-the-art predictors for peptide presentation on MHC class I molecules using an independent dataset, referred to as `MHC-Bench` that we curated from multiple data sources as described in 5.

#### MHC-Bench

The `MHC-Bench` dataset used in evaluating the four predictors consists of 2, 232, 937 peptides of length 9 and 115 MHC alleles. All the MHC alleles in this dataset are human MHC molecules (i.e. Human Leukocyte Antigens or HLA). The `MHC-Bench` dataset contains 3, 464, 013 peptide-MHC pairs previously unseen by predictors during training. It is worth noting that although the peptide-MHC allele combinations were unseen in the training, the peptides by themselves may have been seen in the training data, combined with a different MHC allele. The peptide overlap between training data and peptides across the clusters (Intercluster) for investigated predictors and `MHC-Bench` is shown in Table 1. A description of the construction and composition of the dataset is presented in Section 5.1.

#### MHC class I Predictors

We evaluated the performance of four MHC class I predictors on the test dataset – `MHCflurry`, `MHCfovea`, `NetMHCpan` and `TransPHLA`. The choice of the four predictors was guided by their popularity and performance reported in the literature [11, 29, 38, 43].

***MHCflurry–***2.0 [38] is an ensemble predictor of 10 neural networks for predicting presentation of a peptide. It supports 14, 993 MHC class I alleles. MHCflurry provides three scores, namely – `Binding Affinity` (BA), `Processing score` and `Presentation Score` (PS). PS is produced by a logistic regression model combining the binding affinity and processing scores. A processing score captures the antigen processing probability which when combined with binding affinity substantially improves the performance of the predictor [38].

***NetMHCpan–***4.1 [43] produces `Elution Ligand` (EL) and `Binding Affinity` (BA) score and we refer to them as `NetMHCpan-EL` and `NetMHCpan-BA` respectively. It utilizes NNAlign MA [3], an artificial neural network that predict both BA and EL. For both modes, peptides with rank 2% or less are considered as binders. It supports 11, 000 MHC class *I* alleles.

***MHCfovea*** [29], is an ensemble of multiple CNN models that takes MHC allele sequence and peptide sequence as inputs to predict binding probability. In MHCfovea, ScoreCAM [55] is applied to identify important input positions and residues in MHC allele sequence. It provides the motifs for the first and last 4 positions of peptide for each allele.

***TransPHLA*** [11], is a transformer architecture that predicts binding probability for an input peptide and MHC allele sequence. Using the attention scores, important residues for each position of a 9 *− mer* can be obtain to generate a peptide motif for a given allele.

#### Benchmarking Performance

For determining performance of the MHC class I predictors, we use Area Under ROC (AUROC) and Area Under Precision-Recall Curve (AUPRC) as benchmark metrics. Since peptide binding is MHC allele specific, we calculated performance metrics for each allele separately. The performance metrics for each allele are reported in Supplementary Table 1, 2. The average performance metrics for the MHC class I predictors are reported in Table 2. Across both AUROC and AUPRC, we find all four predictors are comparable in their average performances across all alleles, as seen in Fig. 2c. The predictors achieve a higher performance with the AUROC metric, 0.95 — 0.98, as opposed to AUPRC where it is in the range of 0.75 — 0.86.

**Table 2:** MHC class I mean scores

| Predictor | AUROC | AUPRC |
| --- | --- | --- |
| MHCflurry-PS | $0.983 \pm 0.019$ | $0.858 \pm 0.141$ |
| MHCfovea | $0.979 \pm 0.025$ | $0.795 \pm 0.169$ |
| MHCflurry-BA | $0.980 \pm 0.017$ | $0.794 \pm 0.156$ |
| NetMHCpan-EL | $0.964 \pm 0.043$ | $0.806 \pm 0.158$ |
| NetMHCpan-BA | $0.955 \pm 0.046$ | $0.766 \pm 0.167$ |
| TransPhLA | $0.970 \pm 0.0173$ | $0.752 \pm 0.148$ |

**Figure 2:**
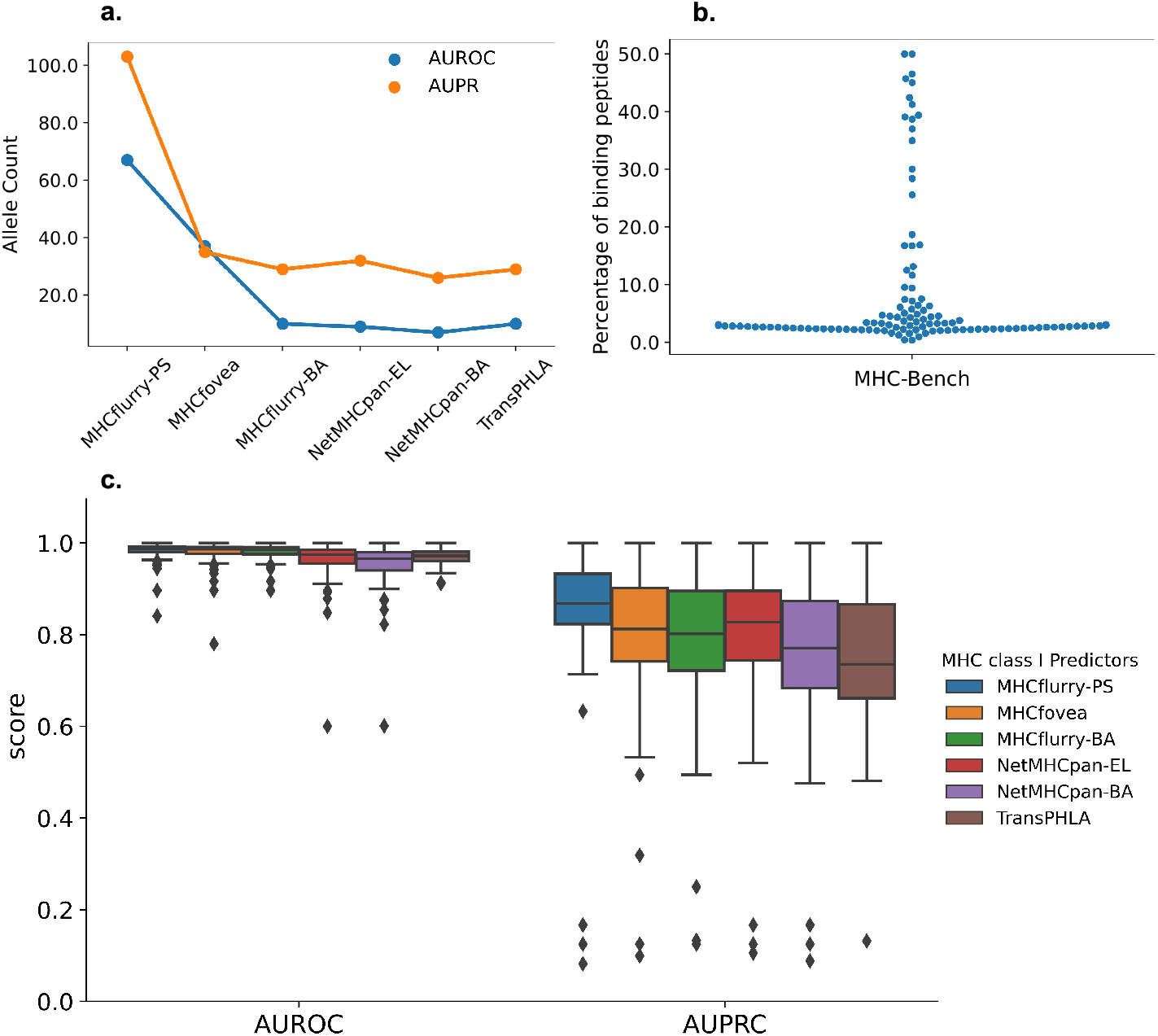
Benchmarking performance of the investigated MHC class I predictors. **a)** Number of alleles for which the predictors are ranked 1 according to AUROC and AUPRC scores. For most of the alleles, MHCflurry-PS ranked 1 followed by MHCfovea. **b)** % binders in `MHC-Bench` dataset per allele. Each dot represents measurement for one allele. **c)** AUROC and AUPRC scores distribution of the MHC class I predictors

For most alleles, there are fewer binding peptides for each MHC allele in comparison to non-binding peptides. This can be seen in the distribution plot in Fig. 2b where for most of the alleles, we only find 1%—10% of the peptides to be binders. In imbalanced scenarios with few positive labels, the AUROC metric can be “overly optimistic” [14]. In contrast, AUPRC is relatively unaffected by this imbalance in the dataset owing to its focus on true positives (because precision and recall do not consider true negatives).

### 2.2 Explanations For MHC Class I Predictors

The explanations for a predictor can be classified as either being *Global* or *Local* explanations. Global explanations for predictor take the form of distribution of input feature values for every output label, observed over *all* inputs in the dataset, thereby, providing an aggregated view of how the model uses input features for a given output label. In contrast, local (or instance-based) explanations, focus on a single input - output instance. Local explanations typically show attribution of input features used for the corresponding output prediction. In the context of MHC class I predictors, binding motifs are examples of global explanations while a vector of attribution values for individual peptide positions forms a local explanation. It is worth noting that our work focuses on post-hoc explanations, i.e., explanations for existing models that have been previously trained. Post-hoc explanations are widely applicable as they can be used over models whose internal structure is not known or is too complex to interpret. Existing MHC class I predictors, like MHCfovea, focus on global explanations by generating binding motifs for MHC alleles. There is limited work on local instance-based explanations for this problem. In next two sections, We motivate the need for local explanations for MHC class I predictors and discuss the additional information it can provide over global explanations.

#### Local Instance-Based Explanations using LIME and SHAP

An instance-based explanations for a 9 *−* mer peptide is a length 9 vector of attribution values, generated using LIME or SHAP. Attribution value of a position can be positive or negative. A positive (or negative) feature attribution value indicates the residue of the position contributes positively (or negatively) to the prediction. Our `MHCXAI` framework allows explanations to be generated for any MHC class I predictor by simply replacing the predictor module while keeping the LIME and SHAP modules unaltered. Using `MHCXAI` framework, we were able to generate LIME and SHAP explanation vectors for input peptides from `MHC-Bench` dataset across all investigated predictors.

For this study, we primarily focus on explanations for peptide input and not for allele input since not all predictors accepts allele input as amino acid sequence. For instance, NetMHCpan and MHCflurry accept allele inputs as `HLA-A`02 : 01 and HLA-A0201 respectively. However, our framework is capable of generating explanations using allele sequences in addition to the peptides, for predictors (such as TransPHLA) that accept allele as a sequence. An example of an allele explanation is demonstrated in Supplementary Fig. S3.

Figure 4a demonstrates examples of LIME and SHAP explanations generated for all investigated predictors for a pair of peptide-MHC allele (`LLVEVLREI`–HLA-A^*∗*^02 : 01). To visualize the explanation, heatmaps are created using the attribution values. The lighter colors indicate positively contribution while darker colors indicate none or negative contribution to the positive class. The peptide `LLVEVLREI` is a binder peptide for allele HLA-A^*∗*^02 : 01 and is correctly classified by all the MHC class I predictors. However, we note that the explanations from SHAP and LIME differ slightly. For instance, in Fig. 4a, both LIME and SHAP place high importance on peptide position P2, but SHAP also recognises peptide position P9 as an important position. P9 would be typically considered an important position for binding as known from literature [9, 45].

**Figure 3:**
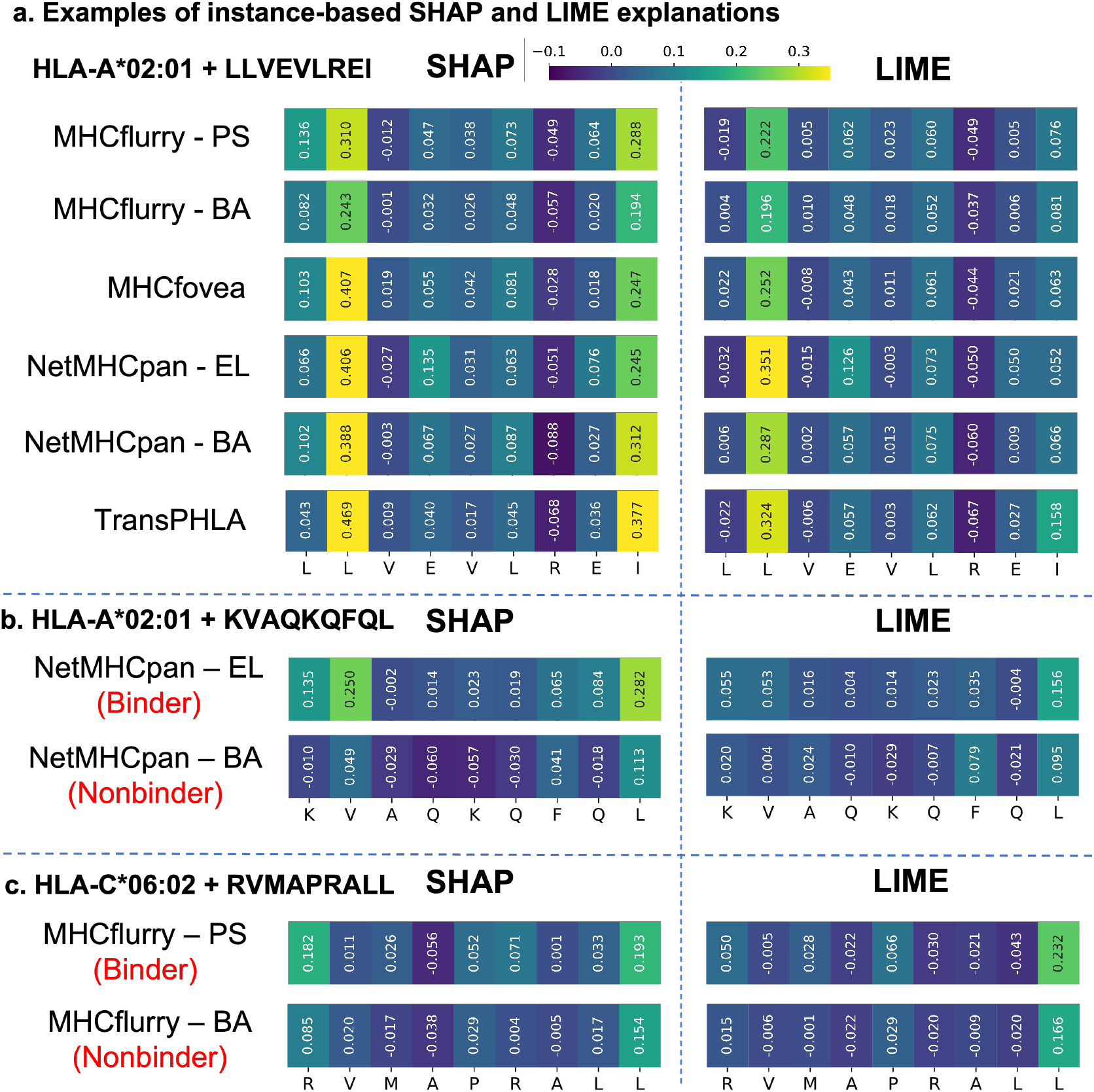
Instance-based (local) explanation for investigated MHC class I predictors. **a)** Examples of SHAP and LIME explanations for all investigate MHC class I predictors for `LLVEVLREI`–HLA-A^*∗*^02 : 01 pair. To visualize the explanations, the attribution values of positions are used to create heatmaps. For each explanation lighter color indicates positive contribution while darker color indicates smaller or negative contribution to the positive class. SHAP explanations for all the predictors highlights peptide positions P2 and P9 as the most important positions for binding while LIME explanations highlight only peptide position P2 as the most important. **b)** LIME and SHAP explanations for NetMHCpan-EL and -BA for peptide `KVAQKQFQL` binding to HLA-A^*∗*^02 : 01. NetMHCpan-EL classifies `KVAQKQFQL` correctly but NetMHCpan-BA does not. SHAP captures this differences in performance and produces different explanations for the two predictor modes. **c)** LIME and SHAP explanations for MHCflurry-PS and -BA for peptide `RVMAPRALL` binding to HLA-C^*∗*^06 : 02. MHCflurry-PS classifies `RVMAPRALL` correctly, but MHCflurry-BA does not. Similar to example in **b**, SHAP captures this difference in performance and produces different explanations for the two predictor modes. In both NetMHCpan and MHCflurry examples, LIME explanations are unable to indicate positions leading to difference in prediction outcome

**Figure 4:**
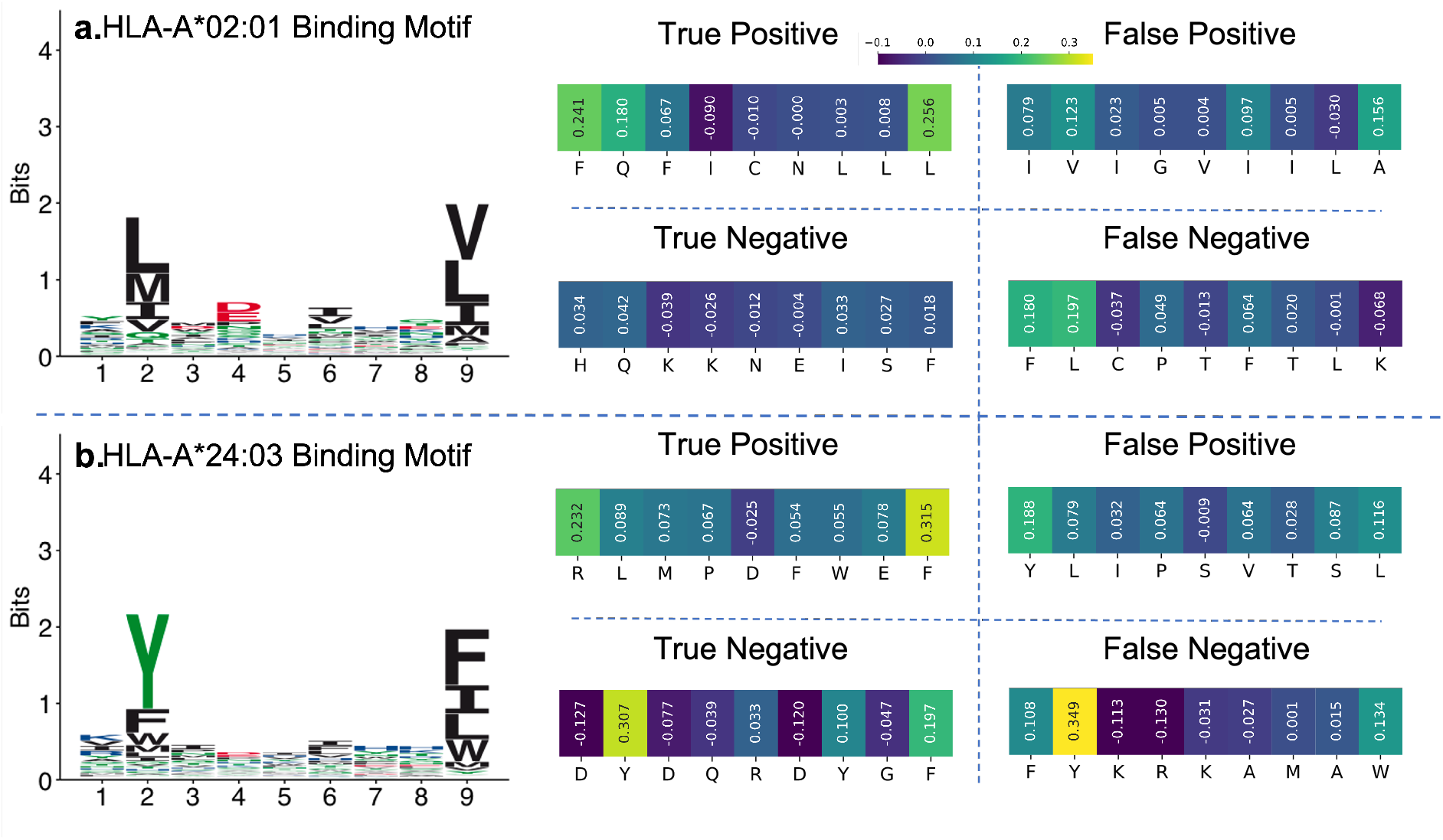
Instance-based (local) explanation for correctly and incorrectly classified peptides along with corresponding MHC allele binding motif. For **a** and **b**, biological binding motifs for HLA-A^*∗*^02 : 01 and HLA-A^*∗*^24 : 03 are obtained from MHC Motif Atlas [53]. For each MHC allele, there are four heatmaps which are SHAP explanations generated for true positive, true negative, false positive and false negative peptides predicted using MHCflurry-PS. The peptides in both **a** and **b** defy the reasoning for binding based on biological motifs. However, the SHAP explanations are able to highlight the cause behind unexpected outcomes.

In Fig. 4b, peptide `KVAQKQFQL` is a binding peptide to MHC allele HLA-A^*∗*^02 : 01. However, within NetMHCpan, NetMHCpan-EL predicts it correctly as binder whereas NetMHCpan-BA predicts it as non-binder. We generated SHAP and LIME explanation for both predictor modes and found that SHAP is able to identify different features (or peptide positions) responsible for the predictions made by NetMHCpan-EL and -BA. For example, for peptide position P1, amino acid `K` contributes positively to prediction outcome in NetMHCpan-EL while it contributes negatively for NetMHCpan-BA prediction. LIME, however, produces similar attribution values for both predictors and is unable to highlight the cause of difference in prediction between the two NetMHCpan modes.

In Fig. 4c, peptide `RVMAPRALL` is a binder to MHC allele HLA-C^*∗*^06 : 02 which is classified as binder by MHCflurry-PS but not by MHCflurry-BA. We explored this difference in prediction using SHAP and LIME explanations. Using SHAP, we identified that for MHCflurry-PS, peptide positions P1 and P9 play an important role. For MHCflurry-BA, peptide position P9 is important position but P1 is not deemed important (See heatmaps in Fig. 4c). This difference in attribution value for peptide positions between the two predictor modes helps isolate the cause for difference in prediction. On the other hand, with LIME, the attribution values in MHCflurry-PS and MHCflurry-BA explanations are similar making it difficult to understand the reason for difference in prediction made by the two MHCflurry modes.

These examples indicate that the XAI techniques can generate explanations for MHC class I predictors, but there is a difference in the explanations generated. Consequently, we assess the validity (using ground truth) and quality (against XAI metrics) of LIME and SHAP explanations in Section 3.

Average time required to generate LIME or SHAP explanation is similar but the time taken to generate explanations for different MHC class I predictors varies widely. For instance, generating one explanation (LIME or SHAP) for MHCfovea takes twice as long as the time taken for generating corresponding explanation for MHCflurry.

### 2.3 Contrasting Local Explanations against Binding Motifs

As stated earlier, global explanations for MHC class I predictors take the form of binding motifs. Binding preference for an MHC allele can be identified by looking at the most commonly occurring amino acids at the anchor positions, which are the primary sites on peptide responsible for peptide binding to an MHC molecule (Position 2 and 9 in a 9*−* mer) [9, 45]. This can be extend to other peptide positions as well and be represented as a binding motif for an MHC allele. Biological binding motifs for an MHC allele are generated using experimentally validated strong binders [42]. In this study, we use biological binding motifs from the MHC Motif Atlas [53] database.

With recent MHC class I predictors, binding motifs are generated using peptides that are predicted as strong binder for an allele. Such peptides are used to generate position specific scoring matrices (PSSM) which are visualized as binding motifs. Binding motifs for MHCflurry, NetMHCpan and MHCfovea are generated in this manner. This motif generated from predictor could be compared against the biological motif to determine if the predictor has learned correct patterns of binding for an allele.

Global explanations, although effective in showing overall trend, tends to miss out deviations observed for certain inputs. Consider, a binding peptide that does not conform to the biological binding motif pattern. For a predictor that correctly classifies this peptide as a binder, it might be worth checking what features the predictor used for the classification. The input features used by the predictor to determine classification for a specific peptide can only be understood with a local explanation, and not a binding motif in this case. Such specialised patterns are difficult to explain with a binding motif.

We illustrate this requirement and the specialized patterns occurring that differ from biological binding motifs with some examples. In Fig. 4a, motif for HLA-A^*∗*^02 : 01 indicates this HLA subtype has a preference for amino acids `L`, `I` and `M` at anchor position P2 which binds to super hydrophobic `B` pocket of the MHC molecule. For such a pocket, water soluble Glutamine i.e. `Q` is an unfavorable amino acid. Yet, solved peptide-bound HLA molecule structures indicate that many peptides with `Q` does bind to HLA-A^*∗*^02 : 01 strongly [37]. An example of such a peptide is `FQFICNLLL` (see Fig. 4a). It is correctly classified as a binder by one of the predictors, MHCflurry-PS. We generated local explanation for this peptide using SHAP and visualized it as heatmaps as seen in Fig. 4a. Highest attribution values were given to peptide positions P1, P2 and P9. High importance of peptide positions P2 and P9 are not surprising as they are anchor positions. However, high attribution value for peptide position P1 makes sense for this peptide binding to HLA-A^*∗*^02 : 01 as position P1 plays an important role in stabilizing the bound structure, as witnessed in [45, 50, 37]. It is worth noting that the amino acid `Q` in position P2 does not appear in the biological binding motif (global explanation) prominently which fails to capture the specialized pattern in this instance. The true negative instance, `HQKKNEISF`, in Fig. 4b also has amino acid `Q` in position P2, like the true positive instance discussed. However, in this case the local explanation shows low attribution values for all peptide positions. This indicates lack of strong binding signal from any of the amino acids in those positions, explaining the negative classification.

Fig. 4b is another example of explanations generated for true positive, true negative, false positive and false negative predictions made by MHCflurry-PS for HLA-A^*∗*^24 : 03. In this example, peptide conforming to binding motif is correctly classified as non-binder whereas peptide not conforming to binding motif is correctly classified as binder.

In summary, instance-based explanations are particularly useful in explaining scenarios where binder peptides do not conform to motifs, misclassifications, and understanding a peptide specific pattern used for prediction. Additionally, we show that global explanations can be created using instance-based SHAP and LIME explanations in Supplementary Fig. S4 which can be useful for quick comparison across models.

## 3 Evaluating XAI Techniques

We consider two popular XAI techniques, SHAP and LIME, and generate an explanation for input peptide as a set of values that indicate the contribution of every peptide position to the prediction output. We evaluate the validity and effectiveness of the explanations with respect to the following metrics,

ΔΔ*G* is difference in free-energy of binding between wild type protein-protein interaction and mutated protein-protein interaction. In mutated protein, a residue in Alanine-scanning mutagenesis the protein is replaced with alanine. This allows identification of residues contributing to the protein-protein interaction which is considered as ground truth for our experiments.

**Consistency** refers to similarity in the explanations produced for two similarly performing predictors on same input.

**Stability** refers to similarity in the explanations produced for two similar inputs (with similar prediction outcomes) for a given predictor.

### 3.1 ΔΔ*G* for Explanation Validity

To trust a predictor, first the explanations generated for the predictors should be trustworthy. A good measure to identify if explanations are good is if they match our prior belief about the input-output relationship [36, 35, 54]. One of the ways to establish this match is by comparing the explanations to the ground truth. In our case, the ground truths are the residues in peptide truly contributing to the binding.

To experimentally identify residues (‘hotspots’) contributing to protein-protein interaction in a bound complex, Alanine-scanning mutagenesis [12] is used which is a resource intensive technique [56]. Computationally, this can be achieved using BAlaS [56, 26] which calculates the difference between the free-We assess consistency of an XAI technique by comparing similarity of explanations for a given peptide between two similarly performing MHC class I predictors Fig. 6a). energy of binding of original bound complex and mutated bound complex where just one residue of ligand peptide is replaced with alanine. This difference in free-energy of binding is indicated as ΔΔ*G* and ΔΔ*G ≥* 4.184*kJ/mol* is considered ‘hot’ or important residue for binding [26]. ΔΔ *G ≤ −* 4.184*kJ/mol* indicates alanine enhances binding relative to the original residue[26]. Any value between denotes neutral substitution [26]. We use this ΔΔ*G* as ground truth for highlighting important residues in peptides.

First, we obtain all the available PDB structures of bound peptide-MHC allele complex listed in MHC Motif Atlas [53]. We then filter the list to bound peptides of length 9 and selected the structures which were correctly classified as binder by all the investigated MHC class I predictors. The final list consists of 250 PDB structures covering 40 MHC alleles (available in Supplementary Table 3).

We compared the ΔΔ*G* with the LIME and SHAP explanations for these peptides. The LIME and SHAP values can be positive or negative similar to ΔΔ*G* which indicates residues contribution to the prediction. We obtained 250 Pearson correlation coefficients corresponding to selected PDB structures for SHAP explanations versus ΔΔ*G*, and LIME explanations versus ΔΔ*G* for each of the investigated MHC class I predictors.

Figure 5a, demonstrates an example of correctly classified peptide `ITDQVPFSV` bound to HLA-A^*∗*^02 : 01. The BAlaS indicates that peptide positions P1, P2, P7 and P9 are ‘hot’ residue (as ΔΔ*G ≥* 4.184*kJ/mol*). The ‘hot’ residues are marked as red in the peptide-MHC complex. The SHAP explanations generated for each of the predictor is shown as heatmap with red arrows indicating the important positions from BAlaS. Overall, we find the models tend to correctly focus on these 4 positions for making the prediction.

**Figure 5:**
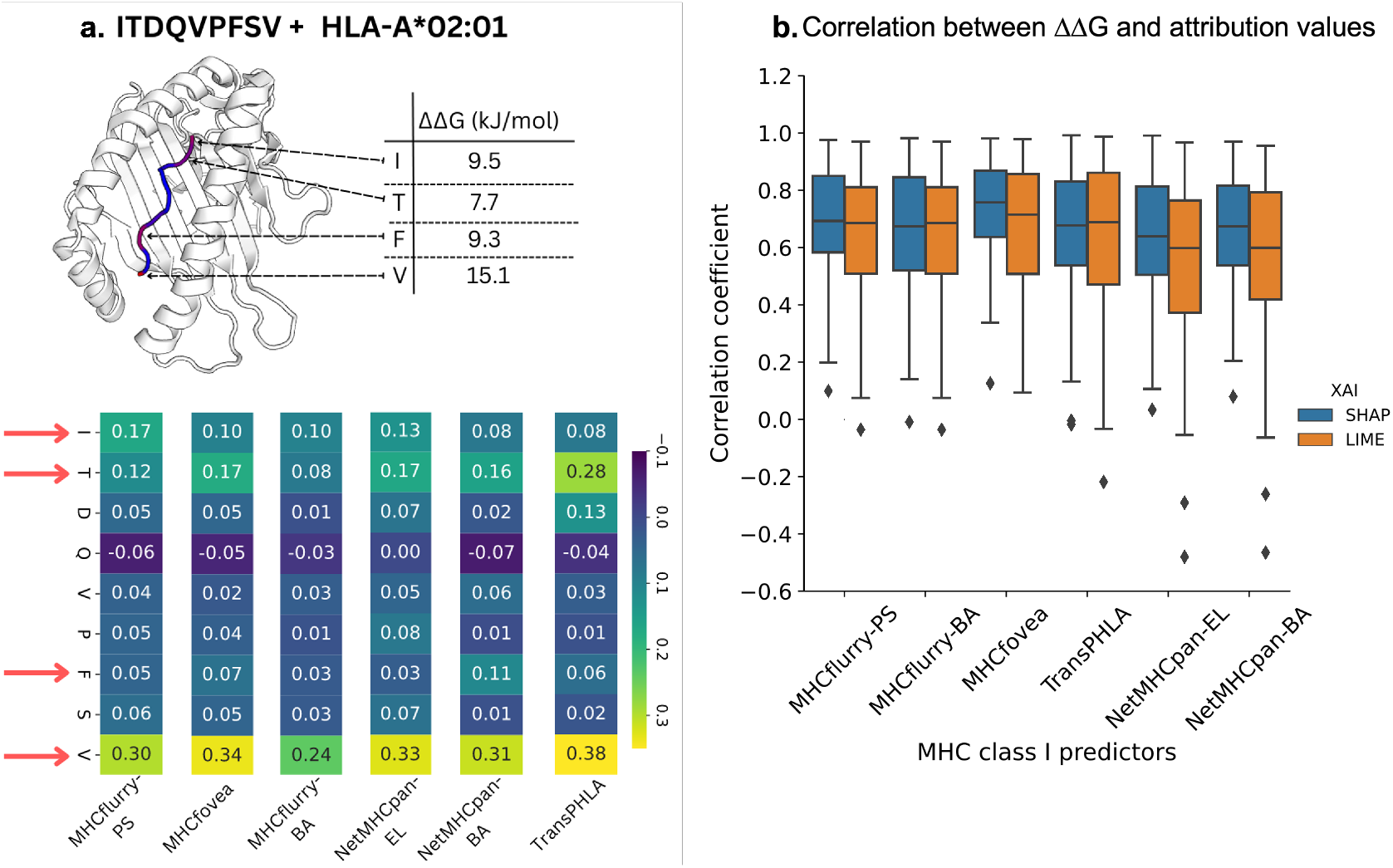
Validation of the explanations is done by comparing the attribution values to the difference in free-energy of binding between wild type protein-protein interaction and mutated protein-protein interaction or ΔΔ*G*. **a)** For `ITDQVPFSV` –HLA-A^*∗*^02 : 01 complex, BAlaS highlights that peptide positions P1, P2, P7 and P9 are very important with respect to binding. Replacement of the residues at these position with alanine leads to an increase in the ΔΔ*G* making the complex unstable. This peptide is correctly classified by all the investigated MHC class I predictors and SHAP explanations are generated for each of them. The explanations mostly match the ground truth as P1, P2, P7 and P9 (Indicated by red arrows) are rightly highlighted as the cause for prediction. **b)** ΔΔ*G* was calculated for each peptide position in 250 PDB structures containing bound peptide-MHC allele complex which were all correctly classified by all the investigated predictors. The SHAP and LIME explanations correlated positively for most complexes indicating that the explanations mostly match ground truth and can be trusted.

However, peptide position P7 with high ΔΔ*G* is not considered important by any of the predictors. This suggest that the other three residues were adequate for the predictors to infer the correct classification outcome. The distribution of correlation coefficient between SHAP – ΔΔ*G* and LIME – ΔΔ*G* (in Fig. 5b) indicates that explanations have positive correlations to ground truth. Overall, we note that the ground truth is more correlated with SHAP explanations compared to LIME explanations.

The variance in the distribution of correlation coefficient observed is expected as BAlaS ΔΔ*G* is only an approximation of the ground truth and suffers certain limitations. One of the limitation is that the ΔΔ*G* calculation is affected by the resolution of the PDB structure (See Supplementary Fig. S6). To mitigate this, we only select PDB structures with highest resolution if multiple structures are available. Additionally, since ΔΔ*G* is calculated by replacing a residue with alanine, the contribution of alanine, if present in the peptide, cannot be ascertained (See Supplementary Fig. S7).

### 3.2 Consistency

We assess consistency of an XAI technique by comparing similarity of explanations for a given peptide between two similarly performing MHC class I predictors Fig. 6a).

**Figure 6:**
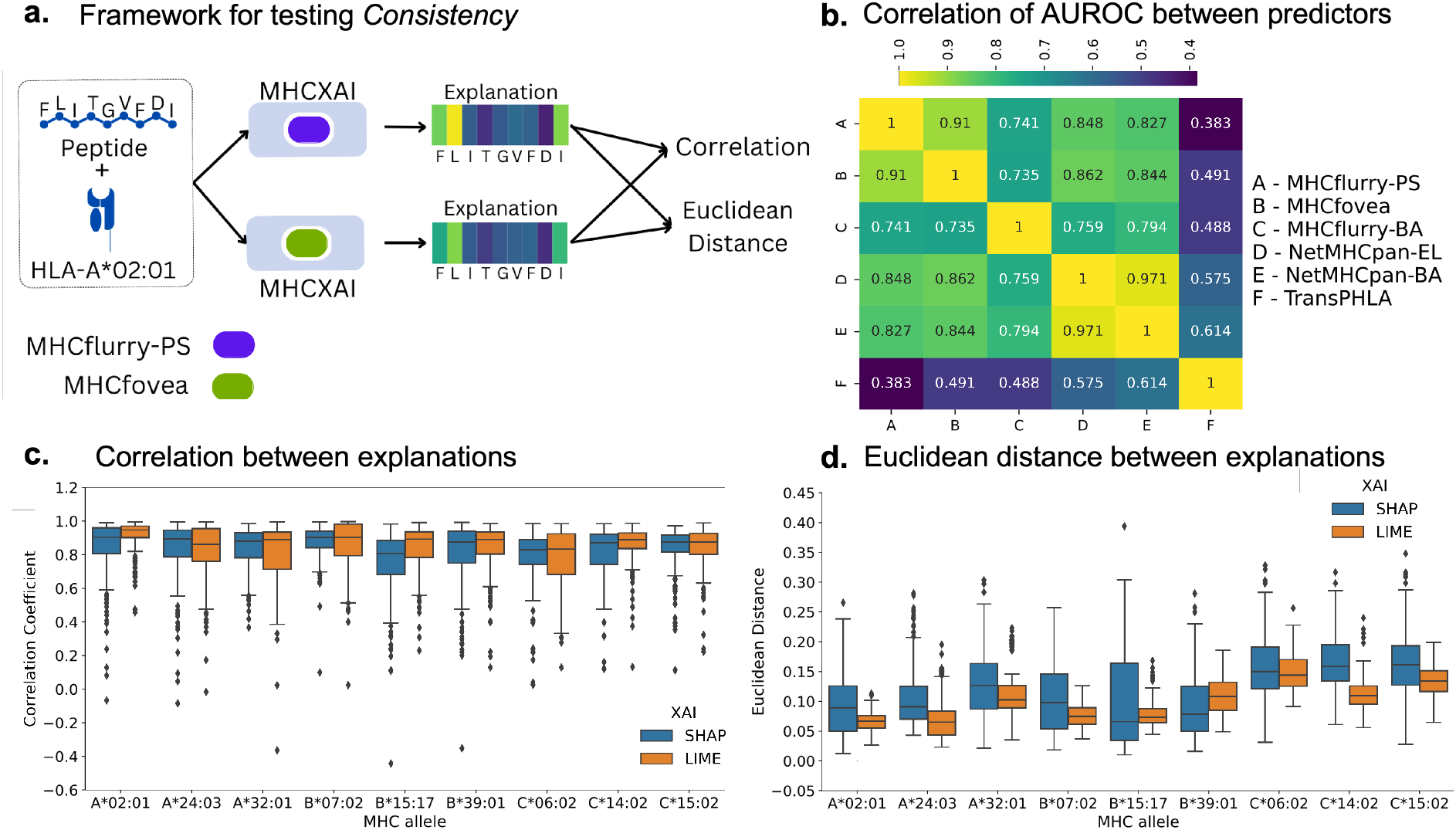
Testing the consistency of the LIME and SHAP explanations. **a)** Illustration of consistency testing framework. **b)** Correlation heatmap for AUROC scores between investigated MHC class I predictors. MHCflurry-PS and MHCfovea are highly correlated in their performances. **c)** For an input peptide, explanations were generated for MHCflurry-PS and MHCfovea using SHAP and LIME. Pearson correlation coefficient was calculated between these explanation and the process was repeated for all 200 input peptides for each of the alleles presented in plot. The distribution of Pearson correlation coefficients is closer to 1 indicating high similarity between the two explanations from two predictors for same input. **d)** In addition to the Pearson correlation coefficient, Euclidean distance was calculated between two explanations from two predictors for same input. For Euclidean distance, values closer to 0 indicate high similarity and high consistency.

To pick two similarly performing predictors, we use the top two predictors from our results in Section 2 (see Fig. 2), namely MHCflurry-PS and MHCfovea. Furthermore, the AUROC scores for the two predictors were highly correlated as reported in Fig. 6b demonstrating the similarity in their performance. To pick two similarly performing predictors, we compared the performance of all the predictors and found that the AUROC scores of MHCflurry-PS and MHCfovea were highly correlated (Fig. 6b). Additionally, these two predictors are also top two predictors from our results in Section 2.

We selected 9 alleles (3 each from HLA-A, B and C) and for each allele, we randomly selected 200 peptides from our `MHC-Bench` dataset and generated local explanations, independently using each of SHAP and LIME, for MHCfovea and MHCflurry-PS. For either LIME or SHAP, to compare the similarity between the explanations from different predictors for each input peptide, we compute Pearson correlation and Euclidean distance between them.

Fig. 6c, reports distribution of Pearson correlation between explanations generated from MHCflurry-PS and MHCfovea using LIME and SHAP, individually, for each of the 9 allele. Overall, most of the explanations were highly correlated for both SHAP and LIME. Fig. 6d, reports distribution of Euclidean distances between the explanations of the two predictors. The explanations which are similar will have Euclidean distance closer to zero. We note that the Euclidean distance distribution for LIME has narrow range and tends to be closer to zero compared to SHAP. This indicates that LIME produces explanations more consistently compared to SHAP.

As a baseline for evaluating the distance between explanations from MHCflurry-PS and MHCfovea, we produce a random explanation by permuting the attribution values of the original MHCfovea explanation. We then compute the Euclidean distance between this random MHCfovea explanation and MHCflurry-PS explanation which is considered as reference. We produce 100 random MHCfovea explanations for every input peptide and average the distance to a reference MHCflurry-PS explanation to create a baseline distance. We create a distribution of baseline Euclidean distances between random MHCfovea explanations and MHCflurry-PS explanations for all 200 input peptides per allele. We compare the Euclidean distance distribution between original MHCfovea and MHCflurry-PS explanations to this baseline distribution. We confirmed that the two distributions for both SHAP and LIME were statistically different using Kruskal-Wallis test at 5% significance level. It is also worth noting that the Euclidean distance for both LIME and SHAP explanations were smaller than the corresponding average Euclidean distance for the baseline for nearly all the input peptides (99% input peptides).

### 3.3 Stability

Stability of an explanation technique refers to the extent to which explanations for similar inputs (with same output) over a given predictor are close. We use the MHCflurry-PS predictor to asses stability of the LIME and SHAP techniques, independently. To identify input peptides that are similar, we perform clustering over a subset of peptide sequences for HLA-A^*∗*^02 : 01. Using GibbsCluster-2.0 [4, 5], we clustered the peptides into 1—10 number of clusters. The number of clusters that yields highest average Kullback–Leibler Distance (KLD) is considered to be the optimum number of clusters. We found choosing 10 clusters has the highest KLD with cluster size ranging between 700—1, 000 peptides. The plot showing KLD distribution and cluster motifs generated from GibbCluster is provided in the Supplementary Fig. S8. Peptides within a cluster are considered similar to each other.

From each of these clusters, we sample 100 peptides which are binders and generate explanations for them with either LIME or SHAP. We then compute the closeness of explanations as the Euclidean distance between them. We compute Euclidean distance between all peptide pairs for the 100 peptides within each cluster. This is used as the *Intracluster distance* distribution. As a comparison, we also compute the distance between explanations for peptides from different clusters, referred to as *Intercluster distance*. We show results for the top 6 most unrelated cluster pairs – (c2,c5), (c3,c5), (c3,c8), (c5,c6), (c5,c9), (c5,c10), based on the similarity of their position-specific scoring matrix, in Fig. 7.

**Figure 7:**
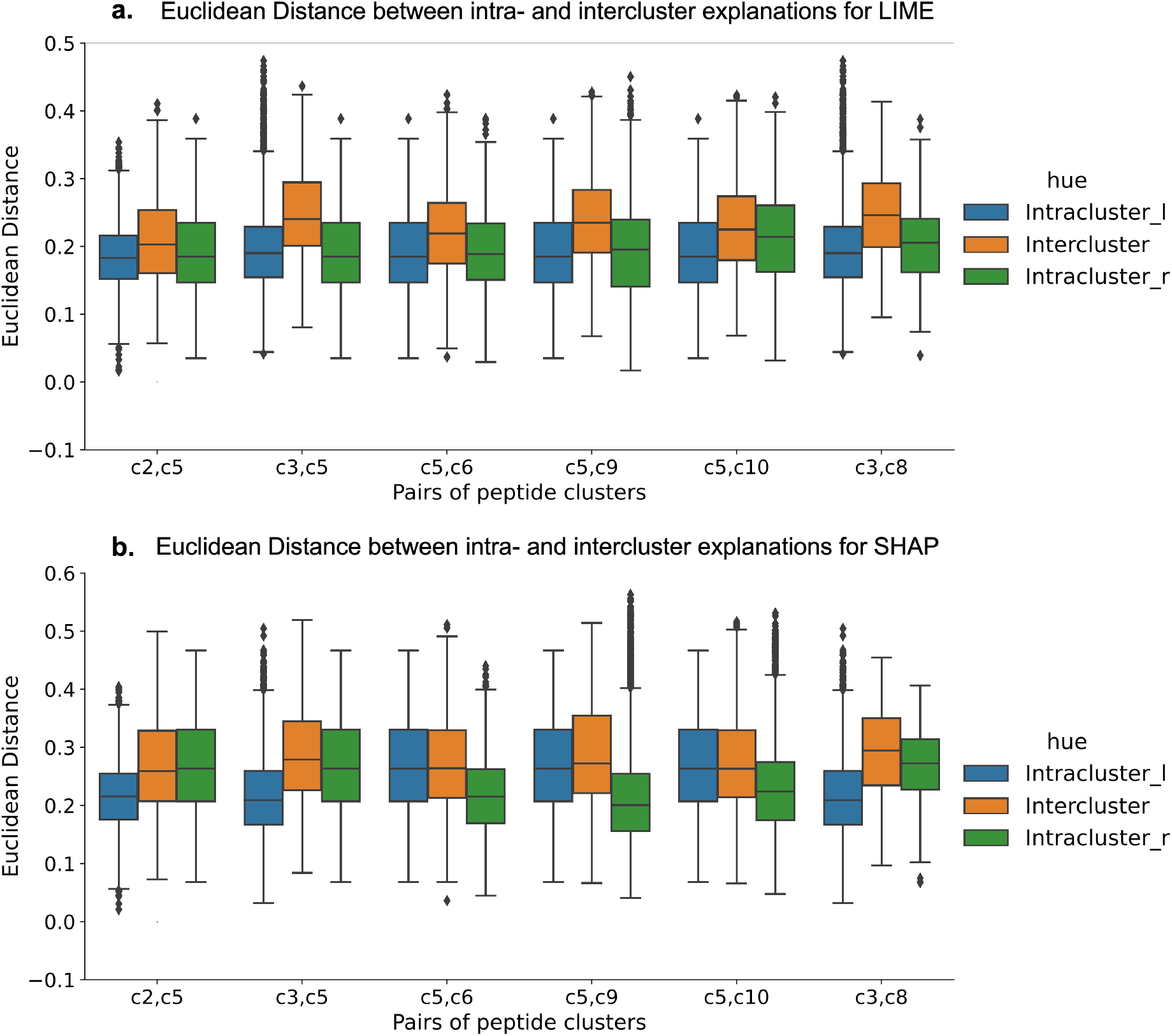
Testing the stability of the LIME and SHAP explanations. Euclidean distance distribution for top 6 cluster pairs (**a** - LIME, **b** - SHAP). For each pair, we have three distributions - `Intracluster_l`, `Intercluster` and `Intracluster_r`. For any two pairs (Say c2,c5), intracluster explanation distance distribution for left (c2) and right clusters (c5) are `Intracluster_l` and `Intracluster_r` while `Intercluster` is distribution of explanations distance between the two clusters

For each pair of clusters in Fig. 7, we have three distribution `Intracluster_l`, `In-tercluster` and `Intracluster_r`. Consider pair `(c2,c5)`, for which `Intracluster_l` is intracluster Euclidean distance distribution for cluster c2 (left cluster), `Intercluster` is intercluster Euclidean distance distribution between c2 and c5 and `Intracluster - r` is intracluster Euclidean distance distribution for cluster c5 (or right cluster). The notation of left-right for Intracluster is arbitrary. We note that intracluster distances are lower than the intercluster distances indicating that LIME and SHAP explanations for peptides within the same cluster is more similar indicating stability. We confirmed the differences between intracluster and intercluster distance distribution in Fig. 7 are statistically significant (Kruskal-Wallis test).

## 4 Discussion

Recently, explainable AI techniques have received a lot of attention as a means of instilling trust in deep learning models by providing interpretability of model decision. Initiatives such as DARPA 2017 XAI research program and EU AI act are geared towards making AI models easy to trust and understand by requiring explanations to help users better understand the decisions made by AI systems. Transparency in AI models is especially important in high-stake scenarios such as biomedical research. In an attempt to bridge the gap between rising importance of XAI and its application in biomedical space, we explored applicability of XAI to MHC class I prediction.

In this study, we used two state-of-the-art XAI techniques namely, SHAP and LIME, to generate explanations for four top performing MHC class I predictors - MHCflurry, MHCfovea, NetMHCpan and TransPHLA. We demonstrate the need for Instance-based or local explanation along with global explanations in fully understanding the predictors. We tested the validity of the explanations by comparing them to ground truth derived from BAlaS of PDB structures of peptide-MHC allele bound complex. We also evaluated the quality of explanations using XAI metrics, consistency and stability. Overall, we found that both LIME and SHAP produce valid explanations which are consistent and stable. While LIME explanations are more stable and consistent than SHAP, SHAP explanations are more accurate.

The presented explanations and their evaluation will help interpret the output of MHC class I predictors and build trust in their decisions. The contributions in this article has scope for generalisation and can be adapted easily to interpret and build trust in other deep learning models over biological sequences.

## 5 Methods

### 5.1 Dataset - MHC-Bench

The `MHC-Bench` dataset is curated by combining multiple existing datasets, namely – Therapeutics Data Commons (TDC) [24, 25], the external and independent dataset from TransPHLA [11], monoallelic benchmark dataset from MHCflurry [38] and benchmark dataset from NetMHCpan [43]. We restricted our dataset to peptides with length 9. We remove peptide-MHC allele combinations that were seen in training data for NetMHCpan–4.1, MHCflurry–2.0, MHCfovea and TransPHLA to ensure fairness in evaluation of the different MHC class I predictors. Additionally, we removed peptide-MHC allele combinations that had conflicting labels and removed MHC alleles where only one class is represented (i.e. either all peptides are binders or non-binders). The final benchmark dataset, named `MHC-Bench`, consisted of 115 MHC alleles and 2, 232, 937 peptides forming 3, 464, 013 peptides-MHC allele combinations. All the MHC alleles were Human leukocyte antigens (HLA).

### 5.2 Explainable AI Methods

For this study, we use SHapley Additive exPlanations (SHAP) [33] and LIME [44] to generate explanations. We created a framework called `MHCXAI` to apply SHAP or LIME for MHC class I predictors.

#### SHAP

Proposed by [33] is a model-agnostic approach which uses a concept from game theory namely Shapley values, to find the contribution of each feature to the model’s output. Starting from one random position mutation, other positions are mutated until the correct model classification is made. This is repeated many times with multiple random perturbation to get the importance of each position, represented as Shapley values. We set number of times the model has to be evaluated to be 25, 000 to reduce the variance in SHAP values (See convergence study in Supplementary Fig. S1. SHAP requires training dataset for each model to generate background distribution for sampling. However, the training data for many of the predictors contains over a million instances which significantly slows the explanation generation process. Instead, we use K-means implementation from SHAP library to summarize the training data as suggested in the documentation. `MHCXAI` accepts peptide and MHC allele as input but only generate samples for peptide from SHAP package as we only focus on peptide explanations in this article. Each of these sample peptides are passed to MHC predictor along with allele to generate predictions. For binding affinity (BA) prediction, the BA values were converted to probabilities by *p*_*BA*_ = 1 *−* log_50,000_ *BA*. The sample peptides and their corresponding predictions are passed to SHAP module to generate explanations for peptides.

#### LIME

LIME [44] is an XAI technique that can be applied to any complex machine learning model such as a neural network (NN). It replaces the complex model locally with something simpler such as linear regression model. LIME creates many perturbations of the original peptide sequence by mutating random positions, and then weights these perturbations by their ‘closeness’ to the original peptide to ensure that drastic perturbations have little impact. It then uses the simpler model to learn the mapping between the perturbations and any change in output label. This process allows LIME to determine which positions are most important to the classification decision. The attribution values generated by LIME can be positive or negative. We use LIME package (https://github.com/marcotcr/lime) for generating explanations. For LIME, the number of samples to generate was set to 25, 000 as high number of samples produces low variance in LIME values. Similar to SHAP, for LIME, `MHCXAI` creates samples only for peptide using LIME package modules. These samples along with allele are passed to MHC predictor to generate predictions. For binding affinity (BA) prediction, the BA values were converted to probabilities by *p*_*BA*_ = 1 *−*log_50,000_ *BA*. The sample peptides and their corresponding predictions are passed to LIME module to generate explanations.

### 5.3 Evaluation of Explanations

#### Validity of the Explanations

To validate the explanations generated by SHAP and LIME, they need to be compared to ground truth. The ground truth in this context are peptide residues contributing the most to the binding to MHC molecule. Experimentally, this is achieved by Alanine-scanning mutagenesis [12] where one-by-one residues in the ligand peptide are mutated to alanine. However, this is resource intensive and can be computationally achieved using BAlaS [56, 26].

We obtained 250 PDB structures covering 40 MHC (HLA) alleles with peptides of length 9 bound to MHC alleles. Using BAlaS, we generated ΔΔ*G* for all 9 peptide positions for all 250 PDB structures. LIME and SHAP explanations were generated for all the investigated predictors for these 250 peptide-MHC allele structures. For each peptide-MHC pair, we correlated the LIME/SHAP explanation vector to corresponding vector of ΔΔ*G*.

#### Explainable AI Metrics

In order to evaluate quality of explanations, various metrics have been proposed [47, 16, 36]. Here, we consider two XAI metrics - `Consistency` and `Stability`.

##### Consistency

This metric captures whether explanations stay the same across similarly performing predictors [36, 47]. Ideally, if models produces similar output, they would be focusing on similar features of data while making a prediction. Therefore, for a model agnostic XAI like LIME or SHAP, we expect swapping the black-box predictor would produce similar explanations for any given input.

##### Stability

This metric assesses extent of similarity of explanations for similar peptide instances [36]. To identify similar peptide instances, we cluster the input peptides for HLA-A^*∗*^02 : 01 using GibbsCluster-2.0 [4], and we expect peptides belonging to the same cluster can be considered similar. GibbsCluster-2.0 is tool which aligns and clusters peptides in an unsupervised manner such that it maximizes the average Kullback-Leibler Distance (KLD) across the clusters. It accepts *λ* parameter ranging from 0 — 1 which represent penalty for intercluster similarity. Since we are interested in clustering binding peptides to a particular MHC molecule, we expect them to be largely homogeneous and we want to be able to detect subtle difference in patterns. To cluster such data, it is recommended to set *λ* to a very low value [4] (*λ* = 0.05). However, this results in overlapping clusters, where the KLD of a peptide within its cluster and across the clusters is similar. Therefore, we select the most unrelated cluster pairs of cluster for testing stability. GibbsCluster-2.0 provides position-specific scoring matrix (PSSM) for each cluster produced using peptides within the cluster. We compare the PSSM of each cluster with other clusters to select the unrelated cluster pairs. In the peptide clusters we sampled 100 binding peptides and generated explanations for them. To assess the similarity of the explanations, we calculate the Euclidean distance between the explanations for intracluster and intercluster peptides. We limit the number of sampled peptides to 100 per cluster as sampling more to generate explanations and calculate distances for all of them would be computational expensive. We expect that, the Euclidean distance for intracluster peptide expanations should be lower than Euclidean distance for intercluster peptide explanations.

## Acknowledgments

We thank Dr. Diego Oyarzun, Dr. Kartic Subr and Dr. Antonio Vergari, Dr. Javier Alfaro at the University of Edinburgh for their valuable comments and feedback. This project has received funding from the European Union’s Horizon 2020 research and innovation programme under grant agreement No. 101017453.

