## Supplementary Text for "Building Trust in Deep Learning-based Immune Response Predictors with Interpretable Explanations"

**Introduction**

The quality of explanation depends on the certain choice of parameters. LIME and SHAP are perturbation based XAI techniques and they produce samples by mutating the original input peptide to evaluate the predictor for generating an explanation. This is one of the parameters that can be passed to the XAI techniques. If the model is not evaluated for an appropriate number of times, the attribution values have high variance. This variance reduces as the number of mutated samples increases. To find the right number of mutated samples to have least variance, we set up an evaluation where we generate explanations for same input peptide by varying the number of mutated samples the XAI technique should produce.

To generate these mutated samples, LIME and SHAP relies on the training data used to train predictor. Here, we explore if there is any difference in explanations generated if we provide training data covering all peptides across alleles or training peptides specific to the MHC allele of interest. We test the validity of the explanations generated by providing these two training datasets.

Furthermore, we show that explanations for alleles can also be generated using LIME [6] and SHAP [5] and global explanations can be formed by aggregation of attribution values. We also demonstrate that LIME and SHAP explanations are mostly correlated and SHAP feature attribution values shows correlation while LIME attribution values are independent.

While BAlaS provides an important alternative to resource intensive Alanine-scanning Mutagenesis [3] to identify peptide residues contributing to binding to MHC molecule, it suffers two major limitations. The first limitation is that the energy calculations BAlaS does over the PDB structure of bound peptide-MHC allele molecule is seemingly affected by the resolution of the PDB structure. Here we demonstrate examples where difference in resolution marks a residue to be contributing to binding in one resolution while not contributing to binding in another resolution. The second major limitation is that BAlaS calculates energies by replacing the residue in peptide with alanine. However, if the input peptide contains alanine residue, the contribution of this residue cannot be calculated by BAlaS.

Finally, we show the GibbsCluster [2] report of clustering input peptides for HLA-A*02:01 used to test stability of the explanations generated by LIME and SHAP.

**Convergence of Explanations**

To generate explanation, LIME produces perturbed samples where it replaces an amino acid with another amino acid sampled from distribution derived from training peptide dataset. Similarly, SHAP evaluates the model by replacing amino acid in test peptide by amino acid from supplied training peptide dataset. As the number of samples generated and evaluated by LIME and SHAP increases, the variance of the attribution value decreases. To test this, we generated explanation for 20 peptides (10 binders, 10 non-binders) for HLA-A*02:01 using LIME and SHAP. We considered explanation generated with 25,000 perturbed samples as baseline. Next, we generated explanations for all 20 peptides using 1,000 – 25,000 perturbed samples and Euclidean distance is calculated from the baseline. In Fig. S1, we see that the Euclidean distance approaches zero for SHAP (Fig. S1a) rapidly as the number of perturbed samples increases until 10,000 samples. While for LIME explanations, Euclidean distance initially drops rapidly until 7,500 perturbed samples, after which the value plateaus and does not approach zero (Fig. S1b). In Supplementary Fig. S1 each dot represents average Euclidean distance calculated for 20 peptides explanation. The errorbar indicates the variance in Euclidean distance and as the number of perturbed samples increases, the errorbar reduces indicating reduction in variance in both LIME and SHAP attribution values.

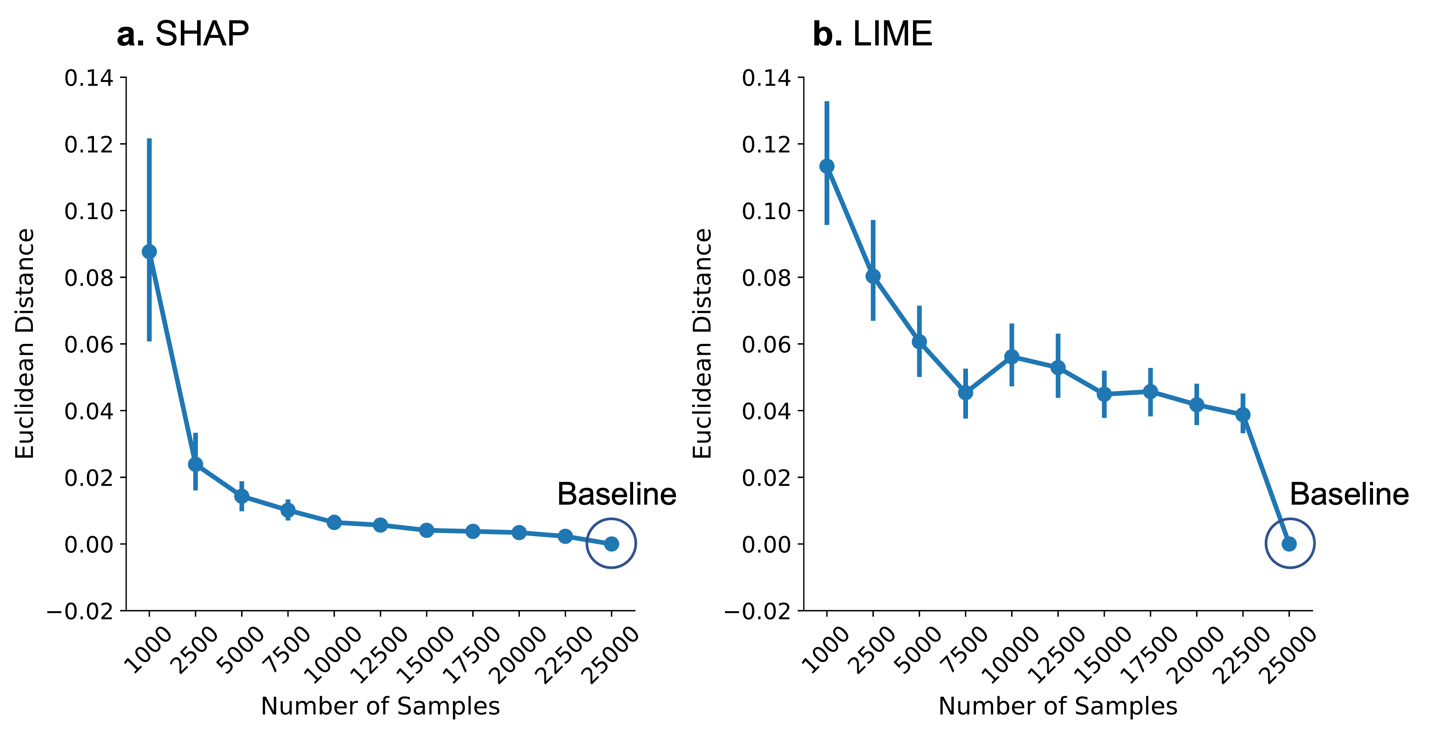

**Supplemental Fig. S1.** Convergence of explanations. LIME and SHAP generates explanations by evaluating model over perturbed samples. The attribution values start to converge from 10,000 samples for SHAP **(a)** and 7500 samples for LIME **(b)**. The SHAP attribution values comes very close to baseline (attribution values generated using 25,000 samples) but LIME attribution values do not eventually converge to baseline. However, as the number of samples increase, the error bar reduces in both SHAP and LIME.

**Impact of Training Data**

Since generating explanations require training data, we also explored if the explanation changes if we provide entire training data across all alleles or just training data specific to the allele of interest. From our PDB dataset of bound peptide-MHC structures, we select structures for alleles HLA-A*02:01, HLA-B*07:02 and HLA-B*35:01. These alleles have many peptides-MHC bound PDB structures (105, 9 and 11 structures for HLA-A*02:01, HLA-B*07:02 and HLA-B*35:01 respectively). We use the ΔΔG for peptide position obtained from BAlaS [4,7] for these structures and generate LIME and SHAP explanations using ‘All training peptides’ dataset and ‘Allele specific peptides’ dataset. For each structure, Pearson correlation coefficient between ΔΔG and explanations is calculated and plotted in Fig. S2a, b for SHAP and LIME respectively. We find that SHAP explanations generated using ‘Allele specific peptides’ are similar to explanations generated using ‘All training peptides’ and are no more or less correlated to the ground truth. On the contrary, we see that limiting training data to allele specific peptides could be detrimental to validity of the LIME explanations as they are less correlated to ground truth than the explanations generated using ‘All training peptides’.

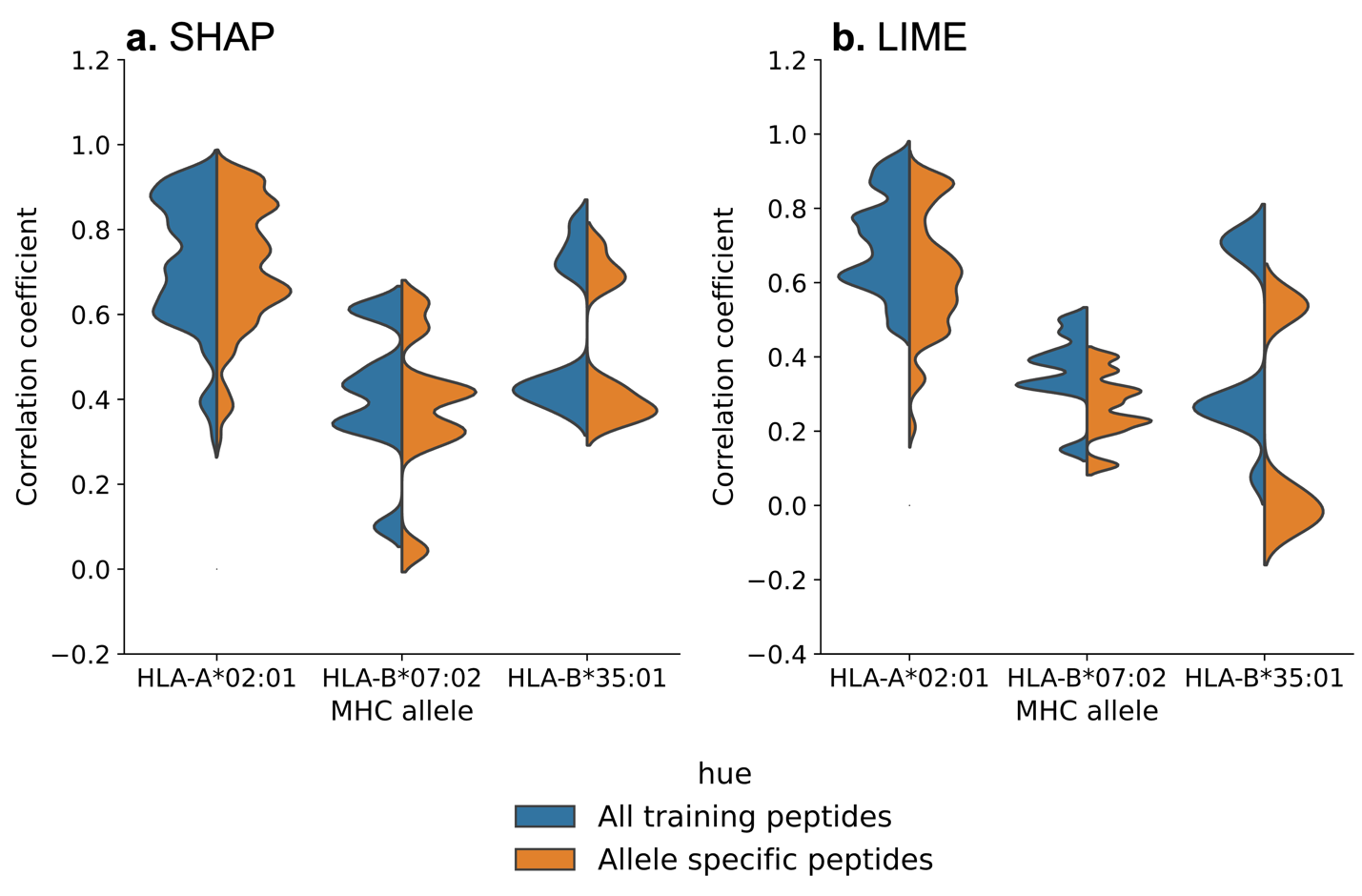

**Supplemental Fig. S2.** Impact of limiting training data on SHAP and LIME explanations. Validity is tested for two sets of explanations generated using ‘All training peptides’ and ‘Allele specific peptides’ against ΔΔG values. **a)** SHAP explanations are not affected either positively or negatively by limiting training data to alleles specific peptides. This is indicated by similar distribution of correlation coefficient between ΔΔG and attribution values generated from two training datasets. **b)** On the contrary, LIME explanations become more unreliable when limiting data to allele specific peptides as seen by difference in correlation coefficient distribution between the two training datasets.

**Explanations for MHC Allele (TransPHLA)**

Similar to peptide explanations, we can also generate explanations for alleles for MHC class I predictor that takes MHC molecule as an input sequence. TransPHLA for example, accepts both peptide and MHC molecule pseudo-sequence as input. A pseudo-sequence of MHC molecule are 34 residues forming binding pocket of MHC molecule’s alpha subunit. To generate explanations for MHC alleles, we use the same framework, but modified it such that the input peptide remains constant, but MHC allele pseudo-sequence is perturbed. In Supplementary Fig. S3, we demonstrate that both LIME and SHAP can generate explanations for HLA-A*02:01 allele sequence for TransPHLA.

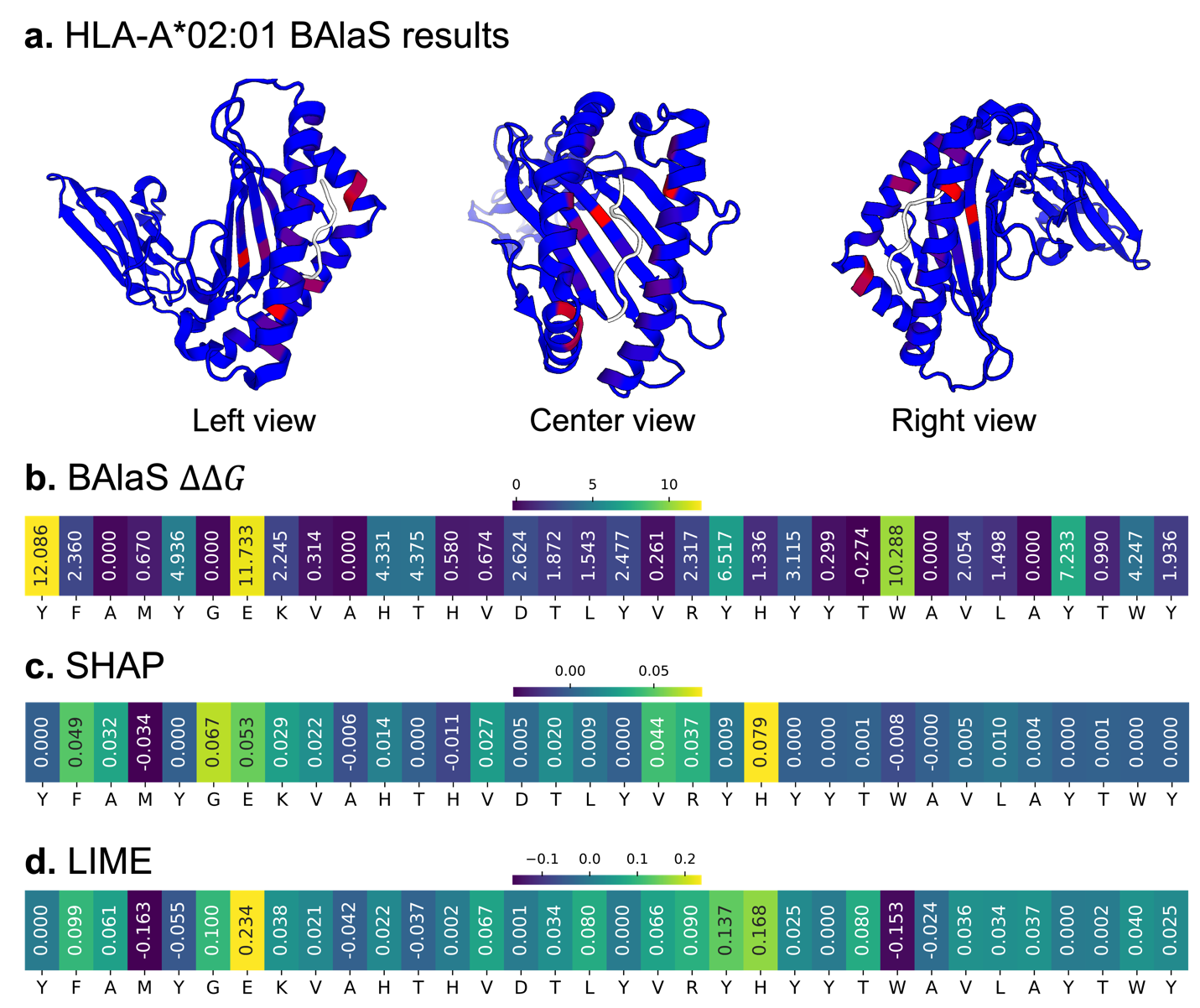

**Supplemental Fig. S3.** Explanation for MHC allele HLA-A*02:01 for TransPHLA. **a)** BAlaS result indicating allele residues contributing to binding (marked in red) from left, center, and right view of the PDB structure. The white molecule is the ligand peptide ‘ITDQVPFSV’. **b-d)** ΔΔG, calculated using BAlaS [4,7], for 34 pseudo-sequence positions of HLA-A*02:01 **(b)** along with SHAP **(c)** and LIME **(d)** explanations for those positions. Explanations for TransPHLA can be generated as it accepts allele input as a sequence of amino acids.

**Global Explanations as Aggregation of Attribution Values**

Local instance-based explanations do not obviate the need for global explanations, they each have their role. For instance, global explanations help understand commonly presented peptide patterns for a tumor that can then be used in a cancer vaccine. It is worth noting that the local instance-based explanations can be aggregated to generate a global explanation. Consider Fig. S4a, c which shows the distribution of SHAP and LIME attribution values for all amino acids at peptide position P1, P2, P8 and P9 for HLA-A*02:01 for MHCflurry-PS. The distribution shown serves as a global explanation across all peptides for MHC allele HLA-A*02:01. The heatmaps in Fig. S4b, d is produced by average all the values from plots like Fig. S4a, c for all positions. We note that the SHAP attribution values for an amino acid have wider range of values whereas LIME attribution values for corresponding amino acid tends to have very narrow range of values. This could explain why LIME produces more stable and consistent explanation than SHAP.

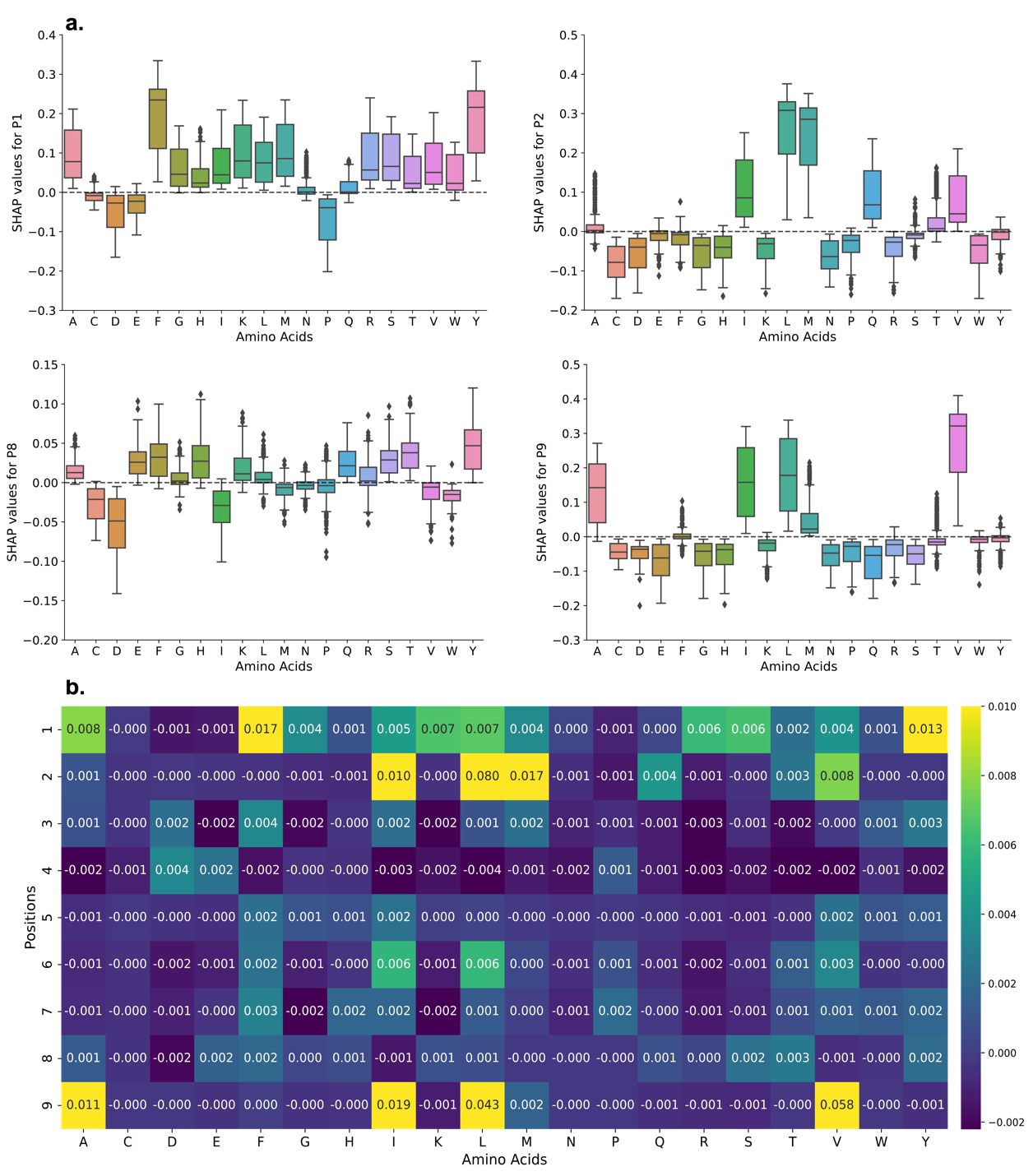

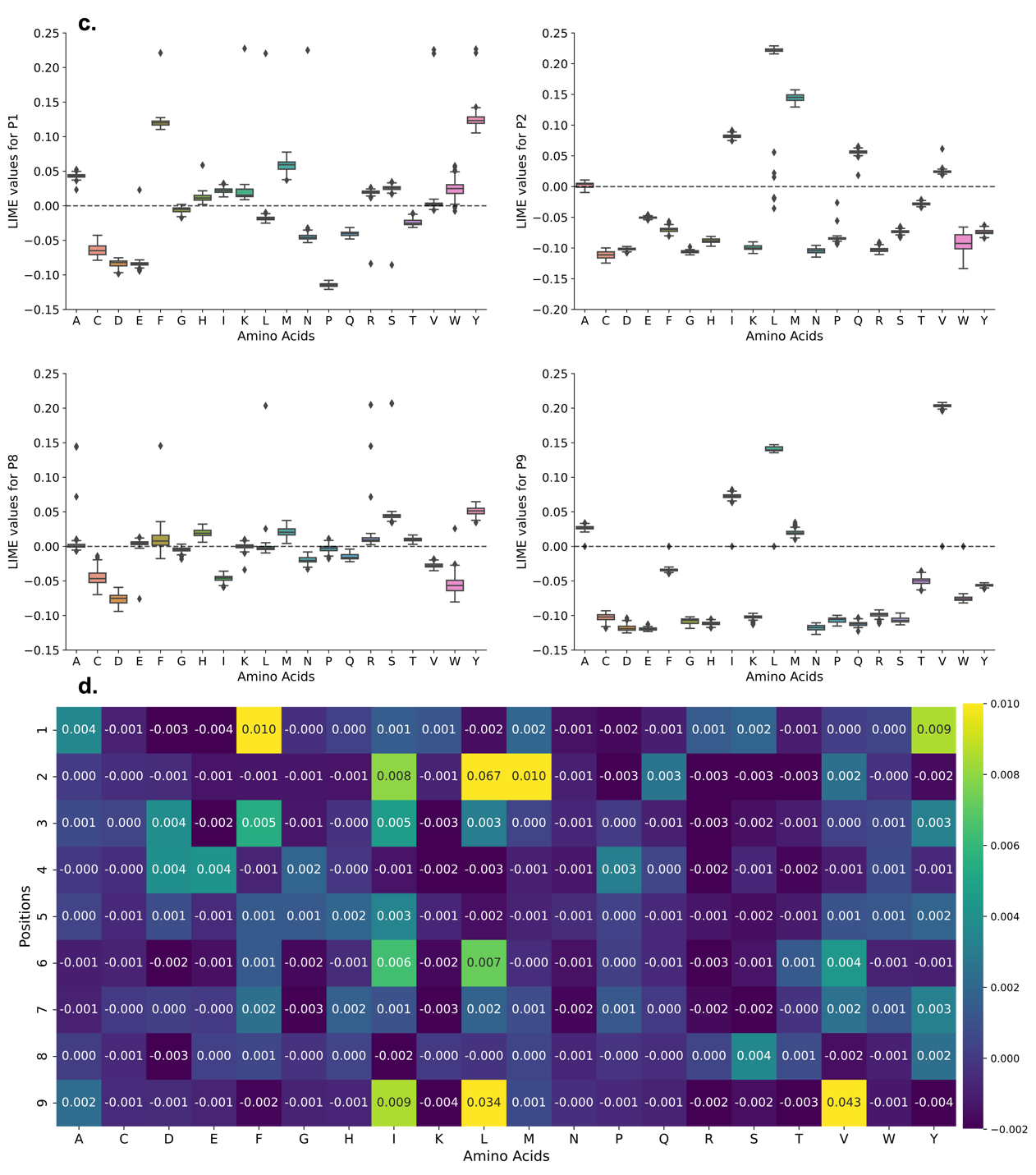

**Supplemental Fig. S4.** Aggregation of SHAP and LIME attribution values to form global explanation. **a)** SHAP attribution values distribution for MHC allele HLA-A*02:01 for all amino acids at N- and C- terminus (P1, P2, P8 and P9) for peptide. **b)** Heatmap of average SHAP values for all amino acids at all peptide positions (P1-P9) for HLA-A*02:01. **c)** LIME attribution values distribution for MHC allele HLA-A*02:01 for all amino acids at N- and C- terminus (P1, P2, P8 and P9) for peptide. **d)** Heatmap of average LIME values for all amino acids at all peptide positions (P1-P9) for HLA-A*02:01. SHAP attribution values for an amino acid tends to have wide distribution compared to corresponding LIME attribution values.

**Properties of Explanations**

LIME and SHAP produces stable and consistent explanations for MHC class I predictors which mostly agree with the ground truth. We wanted to further explore qualities and features of the explanations. First, we explored if the SHAP and LIME explanation for an MHC class I predictor (MHCflurry-PS) agree with each other. Fig. S5a shows distribution of correlation coefficient between LIME and SHAP over 9 MHC allele. We find that the explanations from LIME and SHAP are highly correlated. We also explored if the attribution values in explanations are dependent on each other. Fig. S5b are correlation heatmaps of SHAP/LIME attribution values of peptide positions. For all 3 alleles, SHAP attribution values interdependent (especially for anchor positions P1, P2 and P9) while LIME attribution values were independent. This is possible if SHAP produces test samples from a distribution that has dependent features [1] whereas LIME samples independently for each peptide position from the amino acid frequency generated from training data.

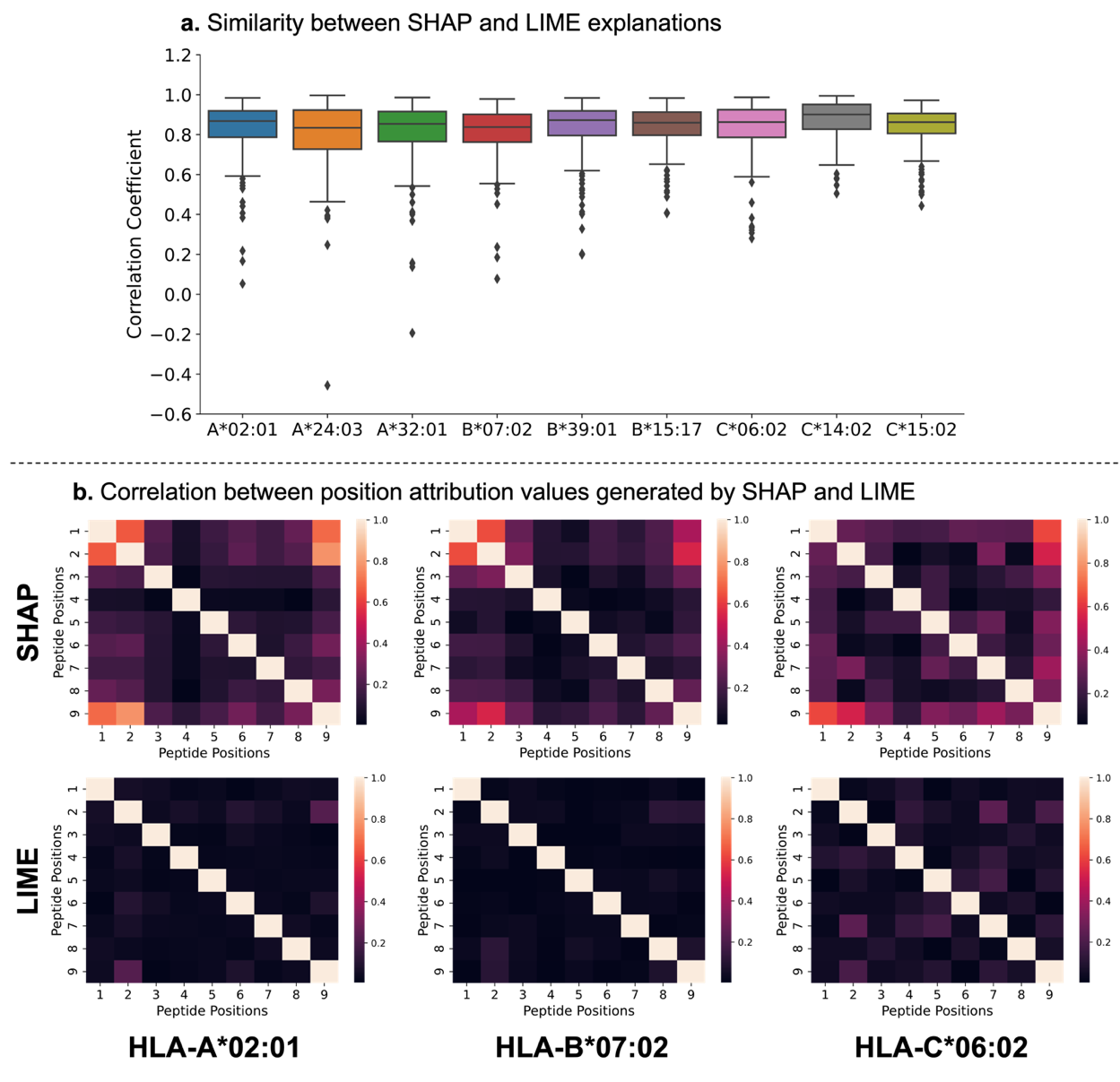

**Supplemental Fig. S5**. Comparison of explanations produced by SHAP and LIME for same input samples and correlation between the attribution values for peptide positions. **a)** The LIME and SHAP explanations are highly correlated for the 9 presented MHC alleles. **b)** Correlation heatmaps for attribution values among all peptide positions indicates that SHAP attribution values can be correlated whereas LIME attribution values tend to be independent.

**Limitation of BAlaS**

BAlaS [4,7] provides an excellent alternative to expensive Alanine-scanning mutagenesis [3] to calculate free-energy of interaction ($\Delta\Delta G$) from PDB structure of peptide ligand bound to an MHC molecule. This $\Delta\Delta G$ can be used to identify peptide residues contributing to binding. However, it suffers two major limitations - $\Delta\Delta G$ calculations are affected by resolution of PDB structure and contribution of alanine residues cannot be calculated. Supplementary Fig. S6 demonstrates the first limitation where for same peptide-MHC allele pair, one PDB structure can cause certain residues to be highlighted as ‘hot’ residues whereas for the other PDB structure it is not. In Supplementary Fig. S6b we see that in 5D2L, residue N has negative $\Delta\Delta G$ whereas for the same peptide-MHC allele pair in 3GSO residue N has neutral effect. The second limitation is seen in Supplementary Fig. S7 where residue at peptide position P2 in both structures is alanine. Here, P2 being anchor residue is appropriately highlighted by SHAP explanation, but BAlaS is unable to quantify the contribution of this position.

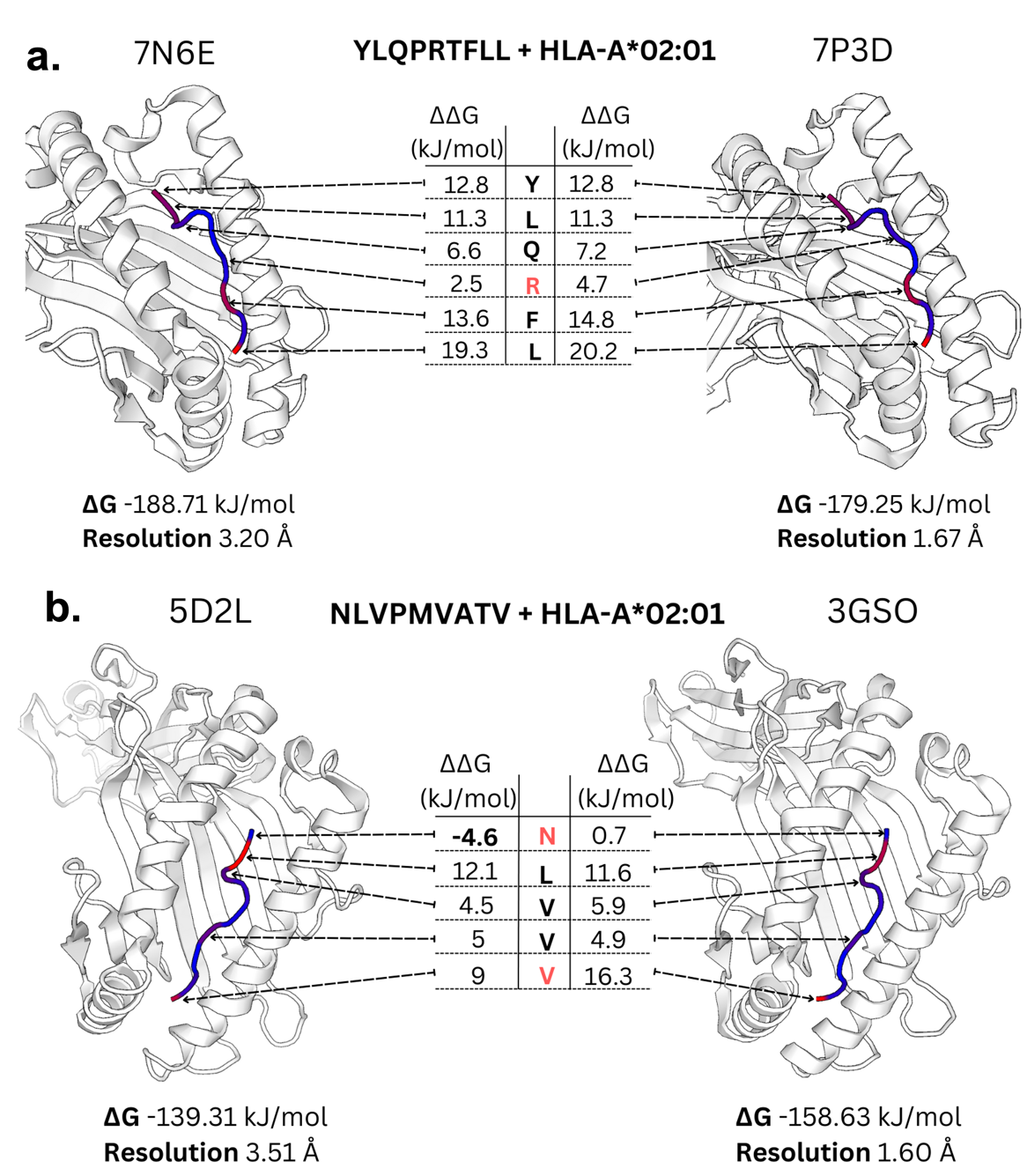

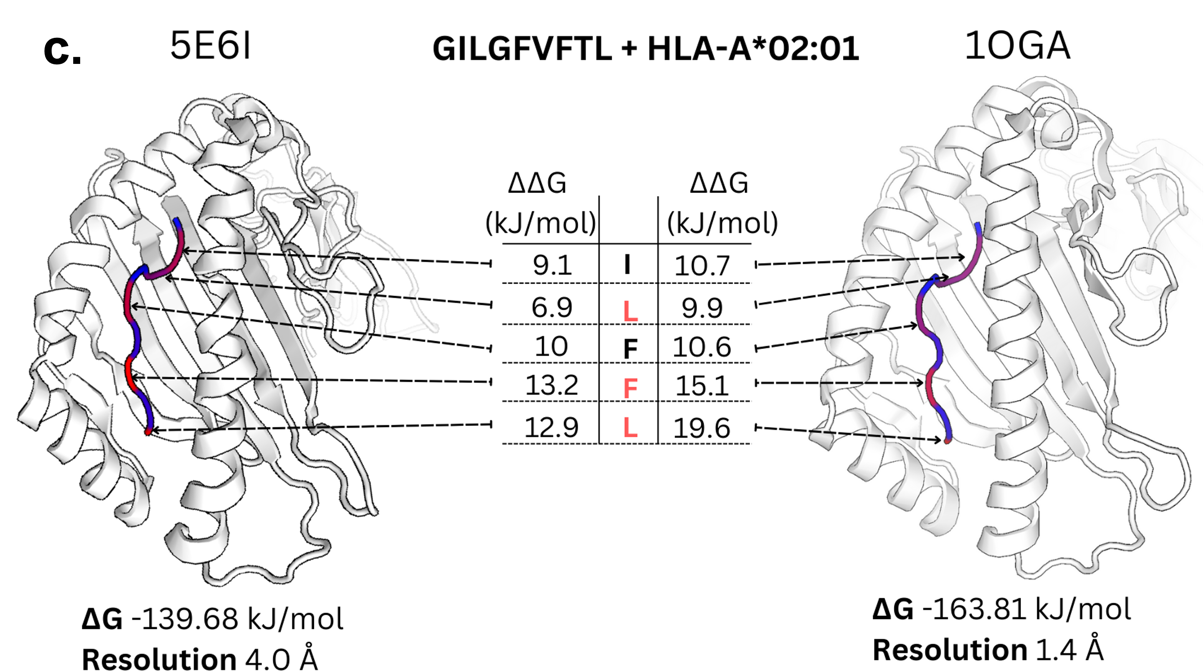

**Supplementary Fig. S6.** Impact of resolution of PDB structure on $\Delta\Delta G$ calculation. In all **a, b** and **c,** there are two PDB structure of different resolutions showing peptide bound to MHC allele. The residues that have relatively large difference in $\Delta\Delta G$ are highlighted in red. The difference in free energy ($\Delta G$) due to difference in resolution can be more than 20 kJ/mol. In **a**, residue R is not highlighted as ‘hot’ residue in coarser resolution PDB structure whereas for 7P3D, $\Delta\Delta G>4.18 kJ/mol$ indicating that it is contributing to binding. In **b,** we see that the residue N is indicated to have negative $\Delta\Delta G$ for PDB structure 5D2L whereas for the finer PDB structure 3GSO, the same residue has no impact.

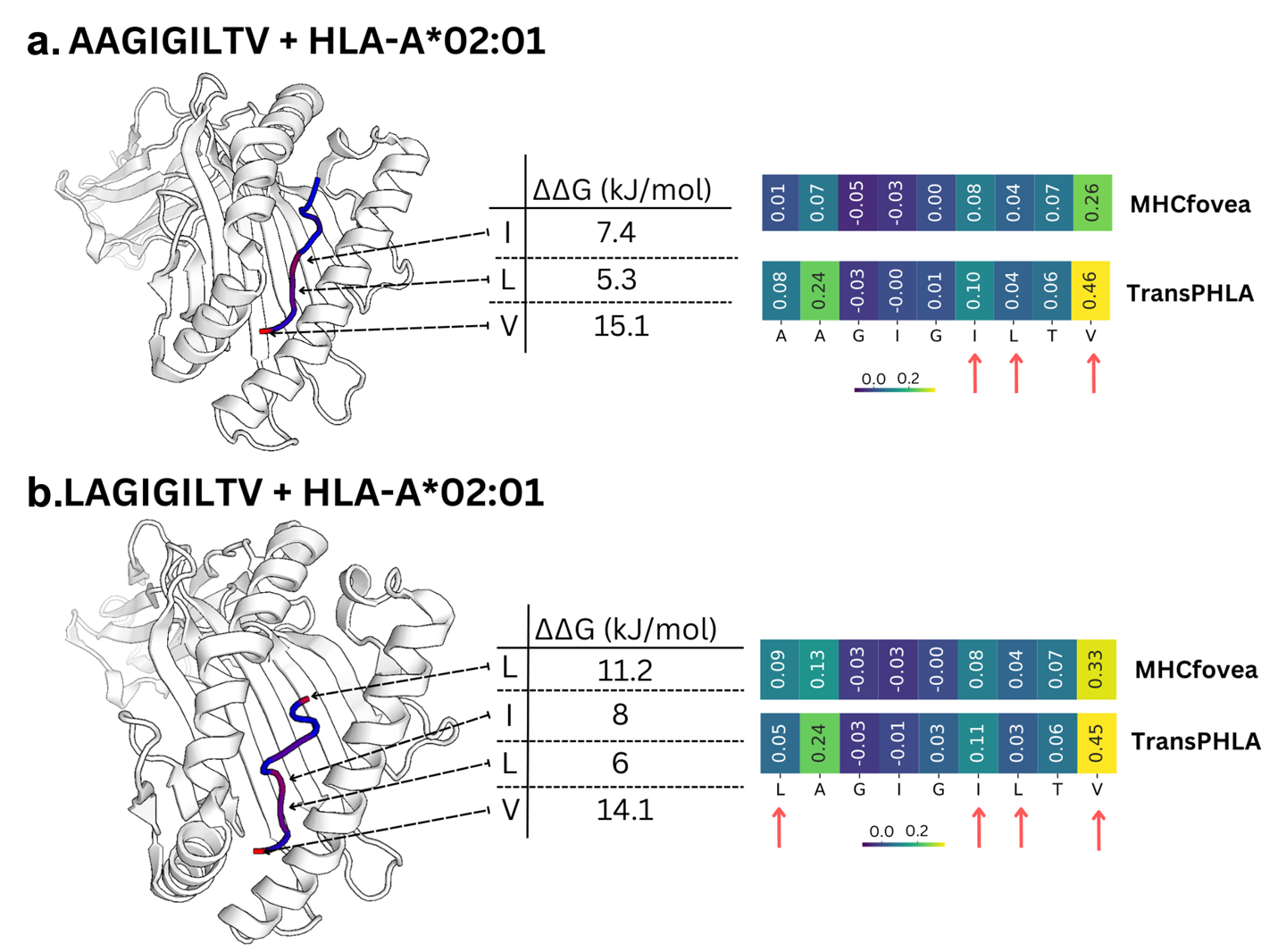

**Supplemental Fig. S7.** Additional examples for comparison between ground truth (ΔΔG) and SHAP explanations and demonstration of limitation of BAlaS [4,7]. Here, peptides ‘AAGIGILTV’ **(a)** and ‘LAGIGILTV’ **(b)** are both binding peptides to MHC allele HLA-A*02:01 with only first amino acid difference. **a)** The BAlaS [4,7] identifies that for peptide ‘AAGIGILTV’, peptide positions P6, P7 and P9 are important for the binding. The two heatmaps are SHAP explanations for MHCfovea and TransPHLA which correctly classified the peptide as binder. The explanations reveal that while P9 is important position, they do not place high importance to other important ground truth positions P6 and P7. **b)** For peptide ‘LAGIGILTV’, P1, P6, P7 and P9 are important residues contributing to the binding. The two heatmaps are SHAP explanations for MHCfovea and TransPHLA which correctly classified the peptide as binder. The explanations reveal that while P9 is important residue, P6 and P7 are not considered highly important for classification even though they contribute strongly to binding. We see that in **(b)** ‘L’ has very high (ΔΔG) indicating the importance of the position. However, neither of the models highlights this to be an important position. TransPHLA indicates P2 as important position contributing to binding which is not indicated by BAlaS. This is important as P2 is one of the primary anchor positions. However, as BAlaS replaces amino acids with ‘A’ to calculate (ΔΔG), we cannot calculate the contribution of ‘A’ towards binding (For example at position P2 in ‘LAGIGILTV’).

**Supplemental Figure**

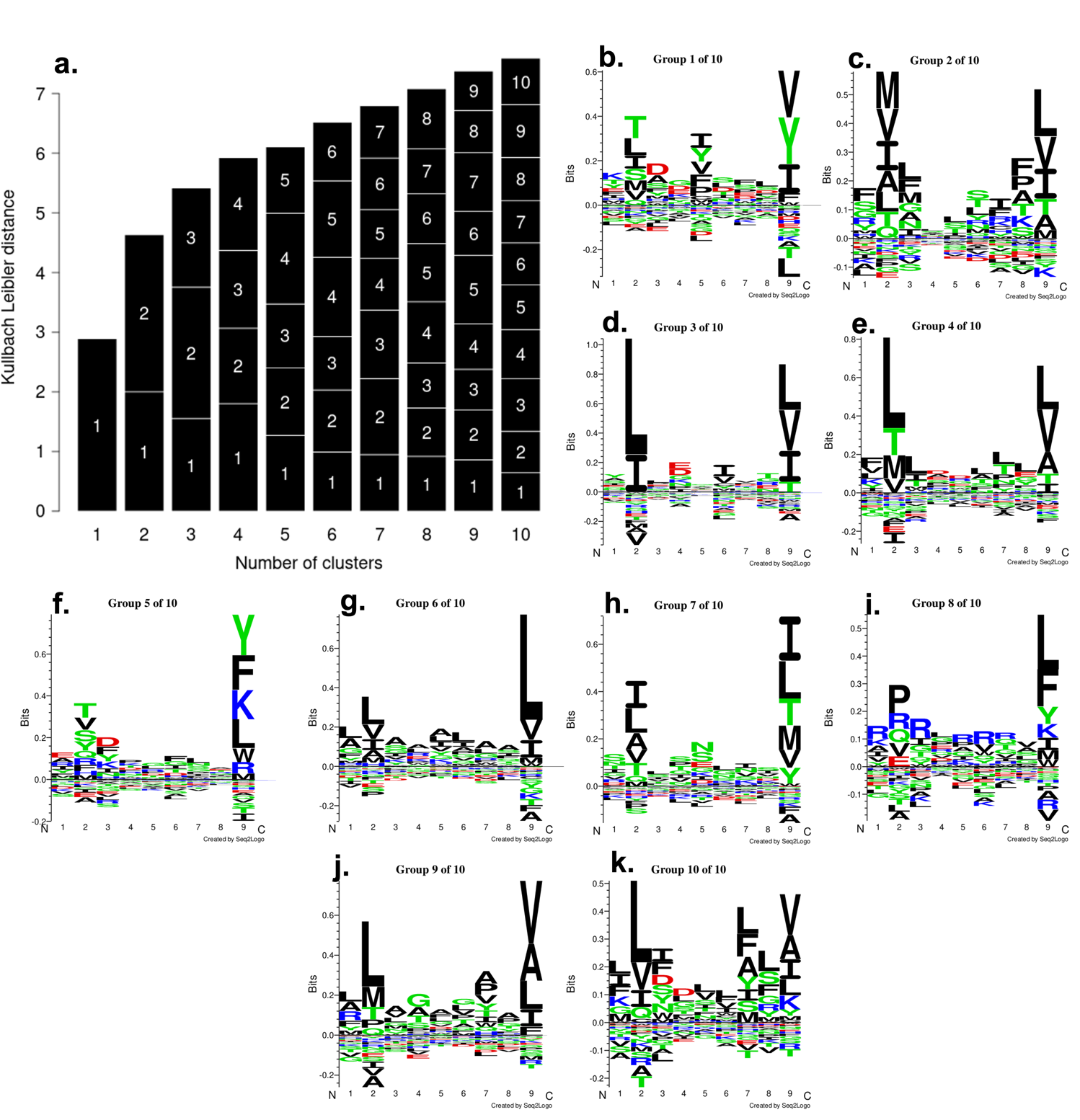

**Supplemental Fig. S8.** GibbsCluster [2] report for MHC allele HLA-A*02:01 peptides. **a)** Kullbach-Leibler Distance (KLD) when peptides are clustered into 1-10 clusters. The KLD increases as the number of cluster increases. The number of clusters with maximum average KLD (Here 10 clusters) is recommended as best clustering result [2] The length of labelled section in each column represents the size of that cluster. For 10 clusters, all the clusters contain nearly the same number of peptides as seen by similar length of sections. **b-k)** The motifs of the peptides present in 10 clusters.
